# Scalp EEG predicts intracranial brain activity in humans

**DOI:** 10.1101/2025.04.07.647612

**Authors:** Ajay K. Subramanian, Austin Talbot, Naryeong Kim, Sara Parmigiani, Christopher C. Cline, Ethan A. Solomon, James Winn Hartford, Yuhao Huang, Ezequiel Mikulan, Flavia Maria Zauli, Piergiorgio d’Orio, Francesco Cardinale, Francesca Mannini, Andrea Pigorini, Corey J. Keller

## Abstract

Inferring deep brain activity from noninvasive scalp recordings remains a fundamental challenge in neuroscience. Here, we analyzed concurrent scalp and intracranial recordings from 1918 electrode contacts across 20 patients affected by drug-resistant epilepsy undergoing intracranial depth electrode monitoring for pre-surgical evaluation to establish predictive relationships between surface and deep brain signals. Using regularized and cross-validated linear regression within subjects, we demonstrate that scalp recordings can predict spontaneous intracranial activity, with accuracy varying by region, depth, and frequency. Low-frequency signals (<12 Hz) were most predictable, with our models explaining approximately 10% of intracranial signal variance across contacts. Prediction accuracy decreased with contact depth, particularly for high-frequency signals. Using Bayesian modeling with leave-one-patient-out cross-validation, we observed generalizable prediction of activity in mesial temporal, prefrontal, and orbitofrontal cortices, explaining 10-12% of low-frequency signal variance. This scalp-to-intracranial mapping derived from spontaneous activity was further validated by its correlation with scalp responses evoked by direct electrical stimulation. These findings support the development of improved inverse models of brain activity and potentially more accurate scalp-based markers of disease and treatment response.

## INTRODUCTION

Traditional approaches to noninvasive brain activity mapping, such as functional magnetic resonance imaging (fMRI), electroencephalography (EEG), and magnetoencephalography (MEG), have provided enormously valuable neuroscientific insights but have significant limitations. Despite its relatively high spatial resolution, fMRI suffers from indirect measurement of neural activity and suboptimal temporal resolution^1,2^. Scalp EEG, despite its relatively high temporal resolution, is limited by poor spatial resolution due to the summative nature of volume conducted signals from many cortical and non-cortical sources that arrive at the scalp^3^; this phenomena particularly obscures the contribution of deep-lying neural sources to the signal measured with scalp EEG. Intracranial EEG (iEEG) recordings, typically used in patients with medication-resistant epilepsy, offer higher spatiotemporal resolution through neural probes but require invasive surgery^4^. Further work is needed to establish a foundation for building noninvasive, generalizable proxies for intracranial activity localized in regions implicated in neurological and psychiatric conditions. This work would have extensive implications in understanding how scalp signals reflect underlying brain activity in perception, cognition, behavior, and pathophysiology.

One way to bridge this gap is through source localization of noninvasive measurements. This methodology consists of 1) establishing an electromagnetic model of the brain, skull, and scalp to model neural signal propagation from hypothetical intracranial sources to the scalp (known as the forward problem) and 2) using the solution for the forward problem as a constraint to posit plausible locations and intensities of intracranial sources that would yield a given profile of recorded scalp activity (known as the inverse problem)^5,6^. While this approach has shown promise in localizing distinct events such as seizures and inter-ictal discharges^7,8^, three critical challenges limit its broader application, particularly to spontaneous activity. First, the fundamental underdetermination of the inverse problem—where multiple distinct neural activity patterns can produce identical scalp recordings—remains a significant challenge for reliably studying neural dynamics during cognition^9,10^, emotion^11,12^, and behavior^13,14^. Second, current methods struggle to accurately localize activity in deeper brain regions^15,16^. Third, and perhaps most crucially, existing approaches to estimate spontaneous activity lack ground-truth validation against intracranial brain recordings^17^.

To address these challenges and develop reliable methods for mapping electrical brain activity from scalp to deeper regions, researchers have simultaneously recorded brain activity from both the scalp and inside the brain^18–21^. While several studies have successfully related scalp EEG or MEG to intracranial brain recordings for epilepsy-related activity^22–26^, to our knowledge only three have explored ongoing brain activity during periods of non-ictal or non-interictal epileptic activity, and only two with EEG^27–29^. However, these two key studies were limited by small sample sizes (N = 3 and 4 patients, respectively) and focused on individual-level analyses rather than cross-patient generalization. Additionally, all three studies never directly evaluated the predictability of the intracranial signal using statistical modeling approaches.

Here, we investigate the predictability of spontaneous intracranial brain activity from simultaneous noninvasive EEG recordings in 1918 contacts in 20 patients with drug-resistant epilepsy. Our study aimed to characterize the relationship between these concurrent recordings via simple and interpretable statistical models. We hypothesized that deep brain activity could be estimated from noninvasive EEG recordings, primarily through its statistical relationship with measurable brain signals from more superficial areas. We expected this estimation to work better for slower brain rhythms, which tend to involve larger brain networks and create stronger, more widespread signals that are easier to detect at the scalp^30–32^. Overall, we observed generalizable, group-level prediction of intracranial spontaneous activity. We found that intracranial activity in the lower frequencies tended to be most predictable noninvasively, with scalp activity in the same frequency band serving as the strongest predictor. At the same time, predictability of neural activity generally declined at higher frequencies. We also observed that the predictability of intracranial signals generally decreased with increased depth of signal. This effect was most prominent when predicting high frequencies. Crucially, we found that prediction of intracranial signals in certain brain regions is generalizable between patients, suggesting a robust relationship between certain types of intracranial source activity and measurements on the scalp, whether by direct volume conduction or indirect co-occurring synchronous activity of shallower (neocortical) sources. Finally, supporting these predictions, we observed that the model scalp loadings for predictability are significantly correlated with evoked scalp maps from direct electrical stimulation to the same intracranial site. Taken together, this comprehensive characterization of scalp-to-intracranial signal relationships provides a foundation for developing noninvasive tools to study deep brain activity in psychiatric conditions, monitor treatment response in neurological disorders, and advance our understanding of the neural mechanisms underlying cognition and behavior.

## RESULTS

### Intracranial activity can be predicted from the scalp with spectral specificity

First, we evaluated in spontaneous recordings whether simultaneously-recorded spectral scalp EEG features can predict spectral iEEG features. For each intracranial contact, we used a time series constructed from computing power in each canonical frequency band in each second of recording from all scalp electrodes to predict time series similarly constructed from that single intracranial site (Fig 1B, see Methods for details). In one example contact in a single patient, we observe greater prediction and predictability in models leveraging low frequency information (Fig 1C-D). This trend of increasing predictability with decreasing frequency was consistent across contacts in that given example patient (Fig 1E-F) as well as across all contacts in all patients (Fig 1H-I, N=20, n=1918 bipolar contacts; mean R^2^ for each participant found in Fig S1). We also observed that, apart from gamma (30-50 Hz) band power, each frequency band in scalp EEG most strongly predicted the same band from intracranial contacts (i.e. scalp theta predicted intracranial theta, scalp alpha predicted intracranial alpha, etc). This appears as higher significantly higher *R*^2^ values along the diagonal of the matrix representing all possible band-to-band comparisons compared to off-diagonal values (Figure 1I, Figure S2). Overall, theta was the most predictable intracranial frequency band on average when considering all input bands, with the highest average R^2^ value across subjects being scalp theta predicting intracranial theta (mean R^2^ of all contacts = 0.09 +- 0.16, N=20, n = 1918; mean R^2^ of *predictable* contacts = 0.32 +- 0.13, N = 18, n = 555; upper quartile R^2^ of *predictable* contacts 0.42 - 0.64). Other most predictive frequency bands included delta (mean R^2^ of all contacts = 0.05 +- 0.13, N=20, n = 1918; mean R^2^ of *predictable* contacts = 0.29 +- 0.15, N = 17, n = 329, upper quartile R^2^ of *predictable* contacts = 0.37-0.71), alpha (mean R^2^ of all contacts = 0.06 +- 0.12, N=20, n = 1918; mean R^2^ of *predictable* contacts = 0.24 +- 0.11, N = 19, n = 515, upper quartile R^2^ of *predictable* contacts = 0.31 - 0.68), and beta (mean R^2^ of all contacts = 0.04 +- 0.13, N = 20, n = 1918; mean R^2^ of *predictable* contacts = 0.29 +- 0.15, N = 17, n = 329, upper quartile R^2^ of *predictable* contacts = 0.29 - 0.76) powers. Gamma and high gamma power were not predictive over all contacts (R^2^ < 0), but, in Gamma, of those regressions that were predictive, they also had the lowest average predictability (mean R^2^ of *predictable* contacts = 0.18 +- 0.11, N = 5, n = 34) (Fig I, Figure S2). Notably, three key trends emerged: 1) low frequency scalp power features were more predictive of intracranial power features across each intracranial band than high frequency scalp power features (Patient 1: Fig. 1G left, Mann-Whitney p = 0.0011, rank biserial correlation effect size rrb = 0.12 ; Group: Fig. 1J left, Mann-Whitney p < 1e-9, effect size rrb = 0.17), 2) low frequency intracranial power features were more predictable than high frequency intracranial power (Patient 1: Fig. 1G middle, p < 1e-9, rrb = 0.35; Group: Fig. 1J middle, p < 1e-9, effect size rrb = 0.23), and 3) diagonal elements of the R^2^ heatmap were stronger than off-diagonal elements (Patient 1: Fig. 1G right, p < 1.4e-8, effect size rrb = 0.20, Group: Fig. 1J right, p < 1e-9, effect size rrb = 0.17). We repeated these tests in both comparing mean R^2^ across all contacts as well as all *predictable* contacts and found these trends remained statistically significant (R^2^ distribution of *predictable* contacts visualized in Figure 1G,J; all test results reported in Table S2). Additionally, we did not observe differences in these spectral predictability trends when grouping patients based on MRI-based epileptic pathology (N = 6 for MRI+ and N = 14 for MRI-; Figure S3). In summary, we found that across our sample, spontaneous intracranial brain activity is predictable from simultaneous scalp activity unrelated to epileptic pathology. Scalp spectral features generally best predicted the same spectral band activity intracranially, with the highest predictive performance occurring with low frequency bands.

**Figure 1:**
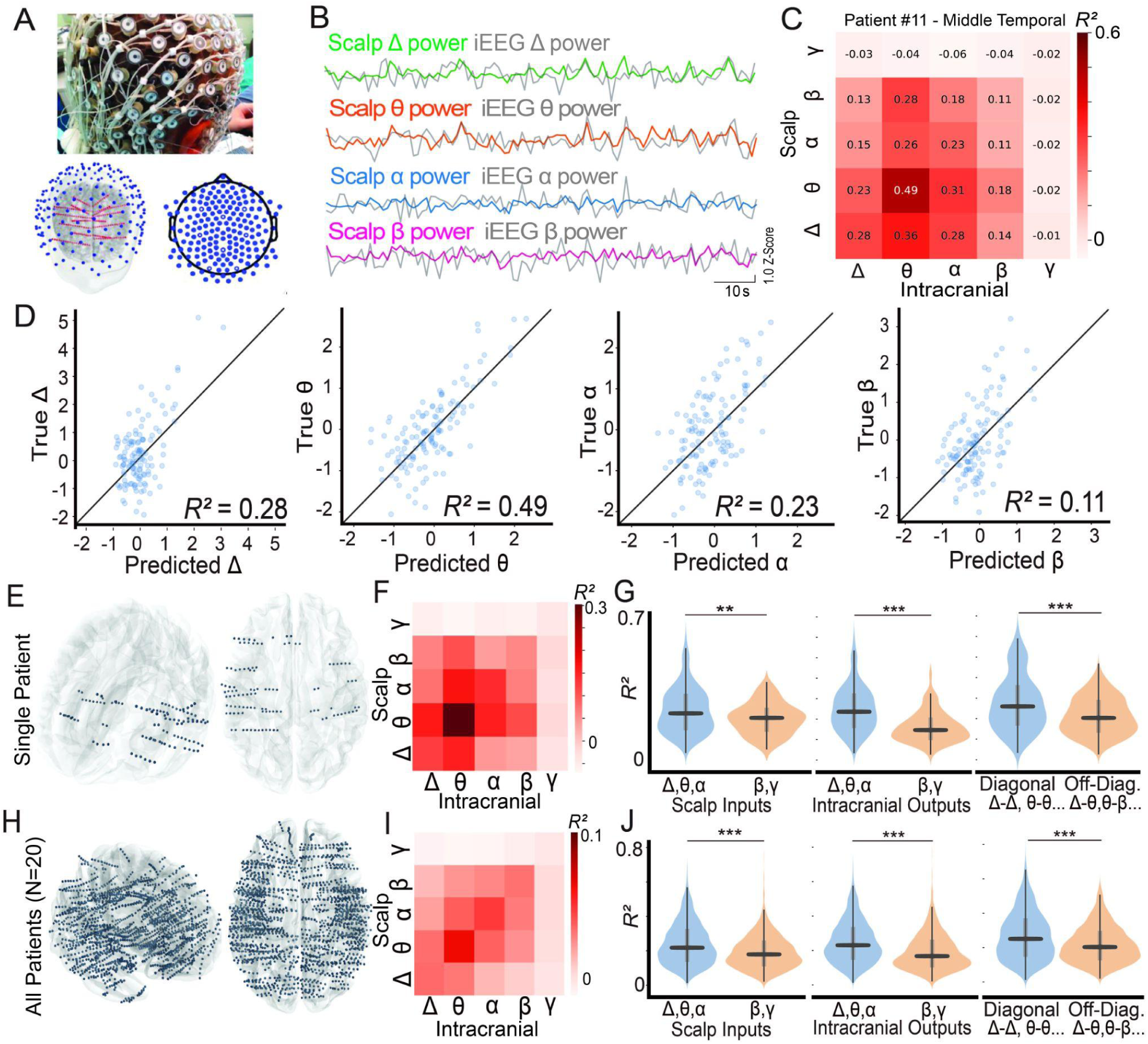
Spectral-specific prediction of intracranial brain activity from scalp EEG recordings. A) Photograph and schematic of patient with simultaneous HD-EEG cap and intracranial stereoEEG contacts. B) Intracranial band power time series overlaid with simultaneous scalp band power time series from a single example intracranial contact. Scalp data is given as the average time series across the total 19 channels. C) Heatmap visualization of R^2^ values of prediction of each intracranial band power time series from scalp frequency band for single example contact. D) Scatter plots of the test-set epochs for four models represented on the diagonal of (C), representing scalp delta predicting intracranial (IC) delta, scalp theta predicting IC theta, scalp alpha predicting IC alpha, and scalp beta predicting IC beta E) Localization of all intracranial contacts in the previous example patient. F) Mean R^2^ heatmap of all contacts in previous example patient. G) Violin plots testing three major hypotheses, visualizing distribution of R^2^ values of predictable contacts from example patient. *left*: First, comparing the mean R^2^ from models using low frequency information from the scalp as input features (scalp delta, theta, and alpha) to those using high frequency information (scalp beta, gamma), Mann-Whitney U Test, **p < 1e-3, ***p < 1e-9; *middle:* second, comparing the mean R^2^ from models predicting low frequency information intracranially (output features - intracranial delta, theta, and alpha) Mann-Whitney U Test, ***p < 1e-9; *right:* third, comparing the mean R^2^ from same-band regression pairs (represented on the diagonal of the heatmap) versus off-band regression pairs (represented by all regressions off the diagonal on the heatmap) Mann-Whitney U Test, ***p < 1e-9. H) Localization of all contacts in all patients. I) Mean R^2^ heatmap across all 20 participants in all contacts (n=1918). J) Violin plots comparing the same three hypotheses as before, but now across all contacts from all patients; Mann-Whitney U Test ***p < 0.001. All statistics, effect sizes, and repeat analysis but for all contacts, including contacts deemed not predictable, are in Table S2.

### The predictability of spectral intracranial patterns is affected by depth and region

Next, we investigated how intracranial predictability is affected by the spatial characteristics of a given recording site. We examined whether specific regions of interest contributed differentially to our finding that scalp EEG best predicted the same frequency band in iEEG (Fig 2a). Our analysis, through evaluating mean predictability in electrodes sorted by Desikan-Kellany parcellation (see Methods), revealed distinct patterns of predictability across brain regions and frequency bands (Fig 2b). Delta band activity was primarily predictable in temporal, insular, and occipital regions, while theta band predictability was more widespread, with strongest predictions in frontal regions (R² = 0.31, 0.32). Alpha band activity showed robust predictability across parietal regions (R² = 0.22-0.25) and frontal areas, particularly pars orbitalis (R² = 0.28). Beta band activity was most strongly predicted in the postcentral parietal cortex (R² = 0.35) and with significant but more modest predictaiblity in supramarginal, superior and inferior parietal regions (R² = 0.21-0.23). Amongst all significantly predictable regions, only middle temporal and lateral occipital regions showed consistent predictability across all frequency bands, and amongst all frequency bands theta exhibited the strongest overall predictability.. Mean R² values of all predictable contacts stratified by region, including those regions not meeting the set threshold coverage of N = 5 patients are reported in Table S3. There were no regions that did not have at least one regression from at least one contact that was predictable, thus all regions are represented in Table S3. The proportion of contacts in a given region that are predictable are reported in Table S4.

**Figure 2:**
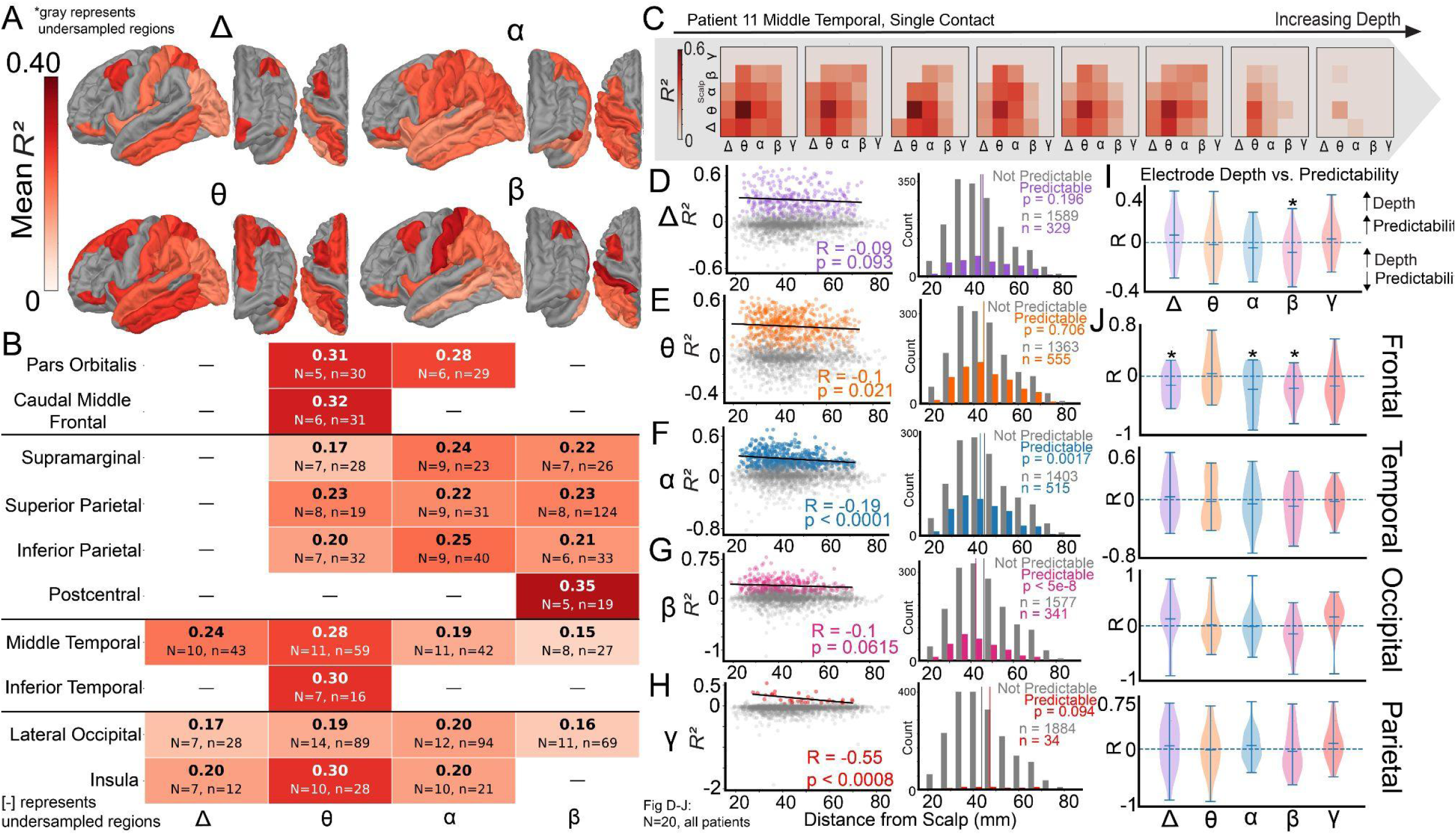
The predictability of spectral intracranial patterns is affected by depth and region. A) Mapping of mean same-band predictability (R^2^) of predictable contacts within a given parcellation, stratified by frequency band. Parcellations with less than 5 participants represented in its population of contacts are excluded (gray). B) Table presenting the mean same-band predictability (R^2^) (shown in A); dashes indicate combinations of region and frequency that did not meet the threshold minimum number of patients to calculate a mean (set to N = 5). C) Illustration of an example single shaft in a single patient that shows depth dependence in spectral predictability. D) Left: scatter and linear regression trend line of all contacts in all patients showing the linear relationship between same-band predictability in the delta band and depth. Right: histograms of predictable contacts (see Methods) within a given band-band pair (in color) and non-predictable contacts (in gray) versus distance. The mean distance of predictable contacts is given by the colored vertical line, while the mean distance of the non-predictable contacts is given by the dark gray line. E-H) Repeat of the analysis of D) for theta, alpha, beta, and gamma. The regression coefficient of distance versus predictability for non-zero contacts is statistically significant for alpha and beta. The difference in mean distance between predictable contacts and unpredictable contacts is statistically significant for delta, alpha and beta (t(14) = -2.3, -2.2, -2.9; p < 0.05 for all) I) Violin plots of the distribution of the correlation coefficients of predictability for all diagonal regressions (delta-delta, theta-theta, alpha-alpha, and beta-beta) versus electrode contact depth for all patients. One-sample t-tests were conducted and distributions with p < 0.05 are shown with a star. J) Violin plots of the distribution of the correlation coefficients of predictability for all diagonal regressions, but stratified by major lobe (See Methods).

The effect of intracranial depth on predictability is of key interest. We hypothesized that as electrode contact depth from the skull increases, predictability from the scalp should decrease. We first illustrate a single-patient example of investigating intracranial predictability versus distance by visually observing a depth dependence on predictability along a single sEEG shaft extending radially out from the deep brain towards the skull (Figure 2C). Across all patients, we assess the effect of depth by examining both *predictable* and *unpredictable* contacts (see Methods for definitions). We first asked whether the depth of predictable contacts was systematically different from unpredictable contacts. To test this, we compared the placement depths of contacts classified as predictable versus unpredictable. For both alpha and beta frequencies, contacts showing predictable relationships between scalp and intracranial measurements were significantly closer to the skull surface than those showing unpredictable relationships. (alpha: mean distance of unpredictable contacts minus mean distance of predictable contacts = 1.9 mm, p = 0.0017; beta: mean distance of unpredictable contacts minus mean distance of predictable contacts = 4.5 mm, p < 5e-8) (Fig 2D-H, Right). However, when examining delta, theta, and gamma frequencies, we found no significant differences in the depth distribution between predictable and unpredictable contacts. (Fig 2D-H, Right, all p>0.05). We then asked, among contacts that were significantly predictable, whether the magnitude of predictability also correlated with depth. We observed a significant negative correlation between predictability and depth for theta, (R = -0.1, n = 555, p = 0.021), alpha (R = -0.19, n = 515, p = < 0.0001), gamma (R = -0.55, n = 34, p < 0.01), and no significant correlation in delta (R = -0.09, n = 329, p = 0.093) or beta (R = -0.1, n = 341, p = 0.062) (Fig 2C-G, Left). Finally, to examine whether these relationships held true within individual subjects, we analyzed how contact depth related to predictability for each patient separately (N = 20, Fig 2I). When evaluating group-level trends, only beta frequency showed a significant negative correlation with depth (p < 0.05), meaning deeper contacts were less predictable. We then investigated whether this depth-predictability relationship varied by brain region by calculating correlation coefficients for each frequency band across four major brain areas (temporal, occipital, frontal, and parietal lobes; Fig 2J). Notably, we observed strong negative correlation in delta, alpha, and beta in frontal contacts (t(14) = -2.3, -2.2, -2.9; p < 0.05 for all). At the same time, the group average relationships in other major regions were not found to be statistically significant (p > 0.05 for all combinations, see Table S5 for statistical test results). In summary, our analysis revealed two key patterns in intracranial EEG predictability: 1) regional specificity, where temporal, parietal, and occipital areas demonstrated consistent predictability across frequency bands, with theta oscillations showing the highest predictability; and 2) depth dependence, where contact predictability decreased with distance from the skull surface, most notably in frontal regions where this negative relationship was significant for delta, alpha, and beta frequencies.

### Spectral signal-to-noise ratio and electrode contact depth have independent effects on predictability

Although certain spatial and spectral trends in intracranial predictability were statistically significant at the group level, there were still multiple confounding factors that could affect the interpretation of these trends. In particular, interpatient experimental variability (as a result of any number of patient-to-patient differences in the recording process, neural activity, or pathology) and interpatient contact placement variability created a need for us to better understand our confidence in the spectral and spatial effects on predictability. Thus, in order to better elucidate the independent effects of our most important variables (interpatient variability, normalized spectral power in the same feature band, electrode contact depth, and absolute spectral power), we chose to fit predictability to a hierarchical Bayesian linear regression model (Figure 3A, see Methods)^33^. As an example of the results of the model, we first observed the mean over all patients of the first two parameters, contact depth and normalized relevant band, for theta-theta predictability, noting the 94% credible interval of the value of the parameter (Fig 3B, C). We then extended this analysis to all spectral combinations of predictability, highlighting the strength and confidence in each effect. A detailed description of each parameter can be found in Methods. We observed that the effect of contact depth is more pronounced when predicting higher frequencies (μ_θ_ = -0.004 < μ_α_= -0.006 < μ_β_= -0.01). Spectral signal-to-noise positively affected predictability across all frequency pairs with a more pronounced effect on theta-theta predictability (μ_θ_ = 0.037). On the other hand, absolute (non-normalized) spectral power values showed small effects on predictability, except for a negative effect on theta-theta predictability (μ_θ_ = -0.018). In summary, our hierarchical Bayesian linear regression model revealed that contact depth reduced predictability at higher frequencies. At the same time, increases in the spectral signal-to-noise ratio led to improvement in predictability (with absolute broadband power having a minimal impact except in theta-theta predictions). These independent effects, established while accounting for inter-patient variability, provide robust group-level insights into the key factors governing scalp-to-intracranial signal relationships, advancing our understanding of how contact placement and signal characteristics influence cross-modal predictability.

**Figure 3:**
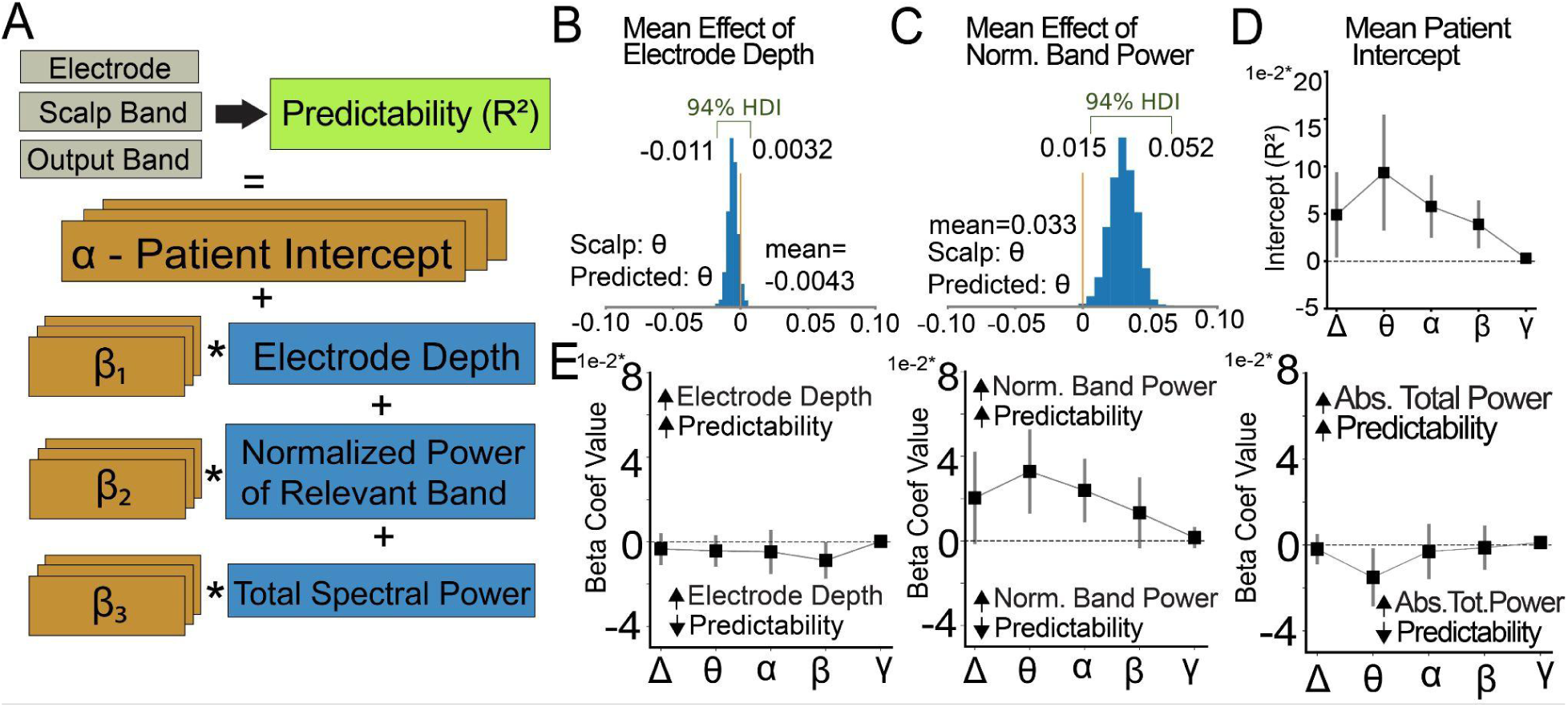
Spectral signal-to-noise ratio and electrode contact depth have independent effects on predictability. A) Schematic representation of hierarchical Bayesian linear regression model for determining effects of listed parameters. Interpatient variability is treated as a random intercept, and normalized spectral power in the same feature band, electrode contact depth, and absolute spectral power are modeled as slope factors for each patient. B) Visualization of the posterior distribution across all participants of the mean regression coefficient associated with contact depth for all contacts in theta-theta regressions. C) Visualization of the posterior distribution across all participants of the mean regression coefficient associated with normalized intracranial band power (in this case theta) for all contacts theta-theta regressions. D) Mean patient intercept for each like-band regression showing that baseline predictability generally varies inversely with frequency E) Plots of each of the model parameters (contact depth, normalized relevant band power, and total spectral power) across increasing diagonal narrowband input/output pairs (theta predicting theta, alpha predicting alpha etc.)

### Intracranial activity from specific brain regions is predictable, and predictions generalize to new patients

To understand how generalizable our predictions of intracranial activity are on new patients, we trained new models focused on three mood-relevant regions—prefrontal cortex (PFC), mesial temporal lobe (MTL), and orbitofrontal cortex (OFC)—using leave-one-out cross-validation (Figure 4A). Models trained in this way explain approximately 10-15% of the variance in intracranial activity in held-out participants (Fig 4B, D, F for OFC, PFC and MTL, respectively, Table S6 for all significant regression values). The strongest predictions were observed between same-frequency bands (e.g., scalp theta predicting intracranial theta in PFC [R² = 0.15, p = 0.03] and MTL [R² = 0.10, p < 0.0001]), particularly in the low-frequency range. Among the three regions, PFC showed the most robust predictability, while significant but more modest predictive performance was observed in the MTL and OFC. We also observe that low-frequency scalp activity (delta and theta) could significantly predict activity across multiple frequency bands in all three brain regions. Additionally, we examined the relationship between the strength of the beta coefficients of each scalp electrode contact compared to the distance away from the intracranial region (Fig 4C, E, G for OFC, PFC, and MTL respectively), with individual scatter plots and trend lines for each set of beta coefficients in Fig. S7. We found strong negative correlations between scalp electrode beta coefficients and their distance from the intracranial region of interest across all three regions, most robust for delta and theta bands (R = -0.47 to -0.75, all p < 0.05). This spatial relationship was also significant for alpha and beta bands in the OFC and PFC but not in the MTL. In summary, our leave-one-out cross-validation demonstrated that scalp EEG can predict 7-15% of variance in regional intracranial activity, particularly in low frequency bands, across PFC, MTL, and OFC regions in held-out patients. The consistency of these negative correlations across multiple frequency bands demonstrates a spatial organization in the cross-modal relationship between scalp EEG and intracranial recordings, with proximal scalp electrodes providing greater predictive power than distal ones.

**Figure 4:**
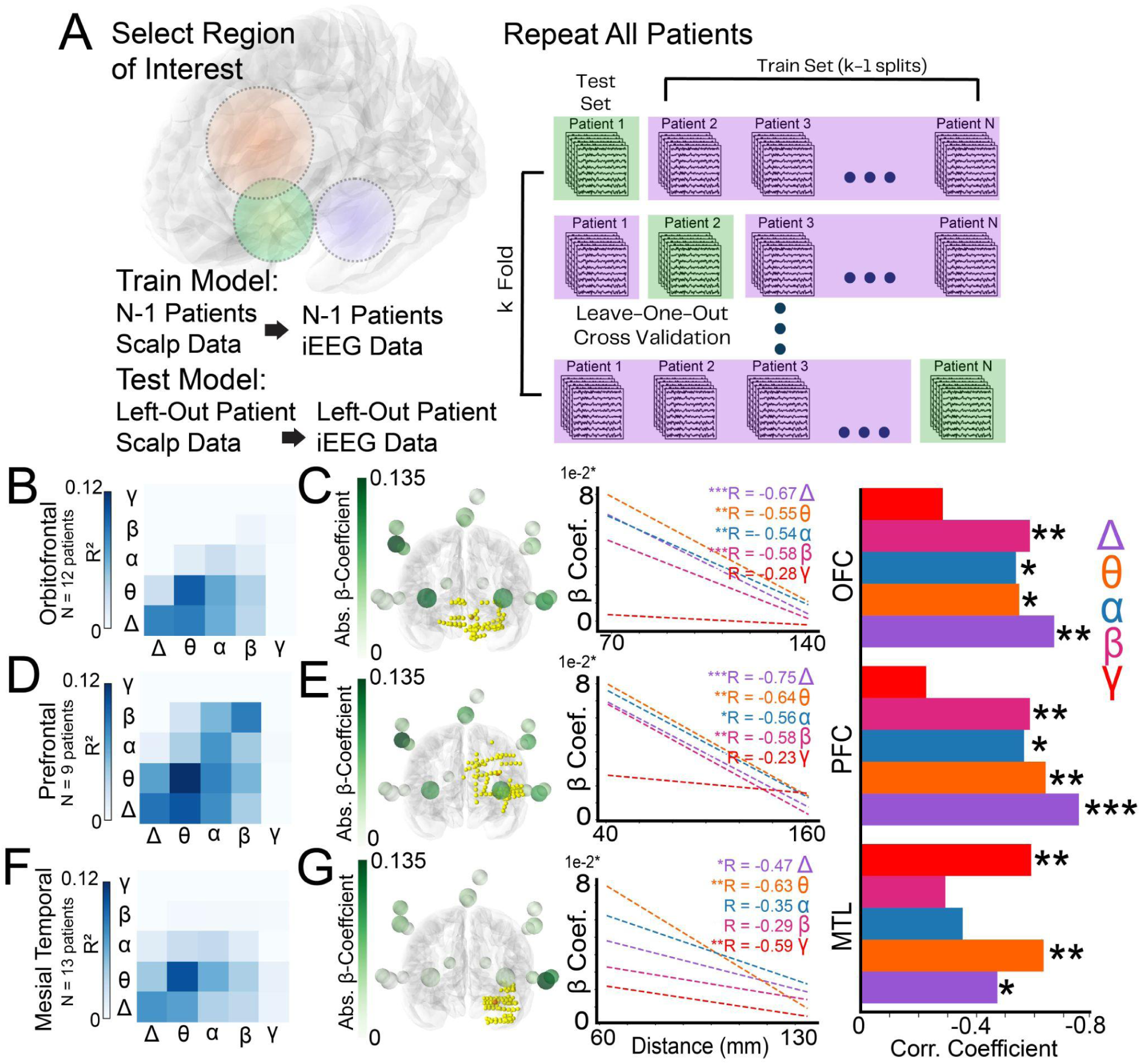
Intracranial activity in mood-relevant regions are predictable and predictions generalize to new patients. A) Schematic representation of the leave-one-out cross validation for each set of regional contact data. B) average left-out-patient predictability for each input feature-output feature pair for contacts located in the orbitofrontal cortex (n=12) C) *left:* First, visualization of average scalp weightings (beta coefficient magnitude) of each of models with theta-theta features. Scalp contacts are colored according to the value of their weightings. All intracranial contacts used in the model are shown. Darker intracranial sphere signifies the region center. *middle*: Trendlines of linear regression models comparing the scalp electrodes weightings of like-band regression models (delta-delta, theta-theta, etc.) versus those scalp electrodes’ distance from centroid coordinate defining the orbitofrontal cortex and *right*: Summary of the R values of each beta coefficient vs. distance trend in the. *, p < 0.05, ** p<0.01, *** p< 0.001 D, F) replicated analysis of B, but for contacts in the Prefrontal region (n = 9) and Mesial Temporal region (n = 13). E, G) replicated analysis of C, but for contacts in the Prefrontal region (n = 9) and Mesial Temporal region (n = 13)

### Direct electrical stimulation correlates with the predictive mapping between spontaneous scalp and intracranial activity

Direct electrical perturbations at specific intracranial sites provide a critical test of our predictive model by revealing functional effective connectivity between these sites and scalp recordings. When stimulation evokes measurable responses at the scalp level, these responses can be compared with our model’s predicted scalp loadings from spontaneous activity. To evaluate this, we leveraged simultaneous scalp and intracranial data during direct electrical stimulation in the same patients. In 11 of the 20 patients studied here, we applied direct electrical stimulation to 83 total intracranial sites. This experimental paradigm allowed us to assess whether the regression model-derived scalp EEG weights from spontaneous intracranial predictions correspond to scalp evoked neural responses induced by focal direct electrical stimulation (Figure 5A, B, C). Direct electrical stimulation of intracranial sites produced temporally distinct components in the scalp-recorded cortico-cortical evoked potentials (sCCEPs^34^; see Figure 5A, B). Following intracranial direct electrical stimulation, we observed early components of the sCCEPs 10-50 ms and late components of the sCCEP 100-200 ms after stimulation (Figure 5A). These temporally distinct early and late components may reflect different underlying neural dynamics, with early responses potentially representing faster processing supported by higher frequency oscillations and late responses corresponding to slower integrative processes mediated by lower frequency bands. We observed that spatial patterns of the sCCEP correlate with prediction weights from resting-state linear mapping models in a frequency-dependent manner. Specifically, beta band (13-30 Hz) resting prediction weights correlated with early components of the sCCEP (R = 0.22, n = 53, p<0.001 versus shuffled data, Figure 5D top), while theta band (4-8 Hz) predictions correlated with late components of the sCCEP (R = 0.19, p<0.001 versus shuffled data, n = 53; Fig. 5D bottom). The strongest correlations overall were seen in the delta (2-4 Hz), theta (4-8 Hz), and beta (12-30 Hz) bands, with resting delta power maps correlating with both early and late components of the sCCEP (R = 0.22 for both), resting theta power prediction correlating with late components of the sCCEP (R = 0.19) and resting beta power prediction correlating with the early components of the sCCEP (R = 0.22) (Figure 5E). All correlations significantly exceeded chance levels based on shuffling analyses (p<0.001; Fig. S8). Theta power prediction weights correlated more strongly with late sCCEP responses compared to early sCCEPs (*p<0.05); conversely, beta power prediction weights correlated more strongly with early sCCEP responses compared to late responses (*p<0.05) (Figure 5E). Finally, we observed some variation in correlation strengths between sCCEPs and resting scalp-to-intracranial predictability by brain region (R = 0.11-0.35). Region-specific frequency preferences emerged, with the cingulate and insula sCCEPs correlating with resting theta predictability (mean R > 0.25 across patients) and lateral temporal regions sCCEPs correlating with resting alpha predictability (R > 0.20 across patients; Fig S9 A, B). The frontal, cingulate, and parietal regions showed significant differences in how their early versus late sCCEP responses correlated with band power predictions (Figure S9C). In summary, direct electrical stimulation experiments demonstrated that our predictive mapping approach is statistically related to stimulation-evoked propagation patterns of electrical activity, with distinct temporal components of scalp-recorded responses showing frequency-specific correlations that vary by brain region across our sample of patients.

**Figure 5.**
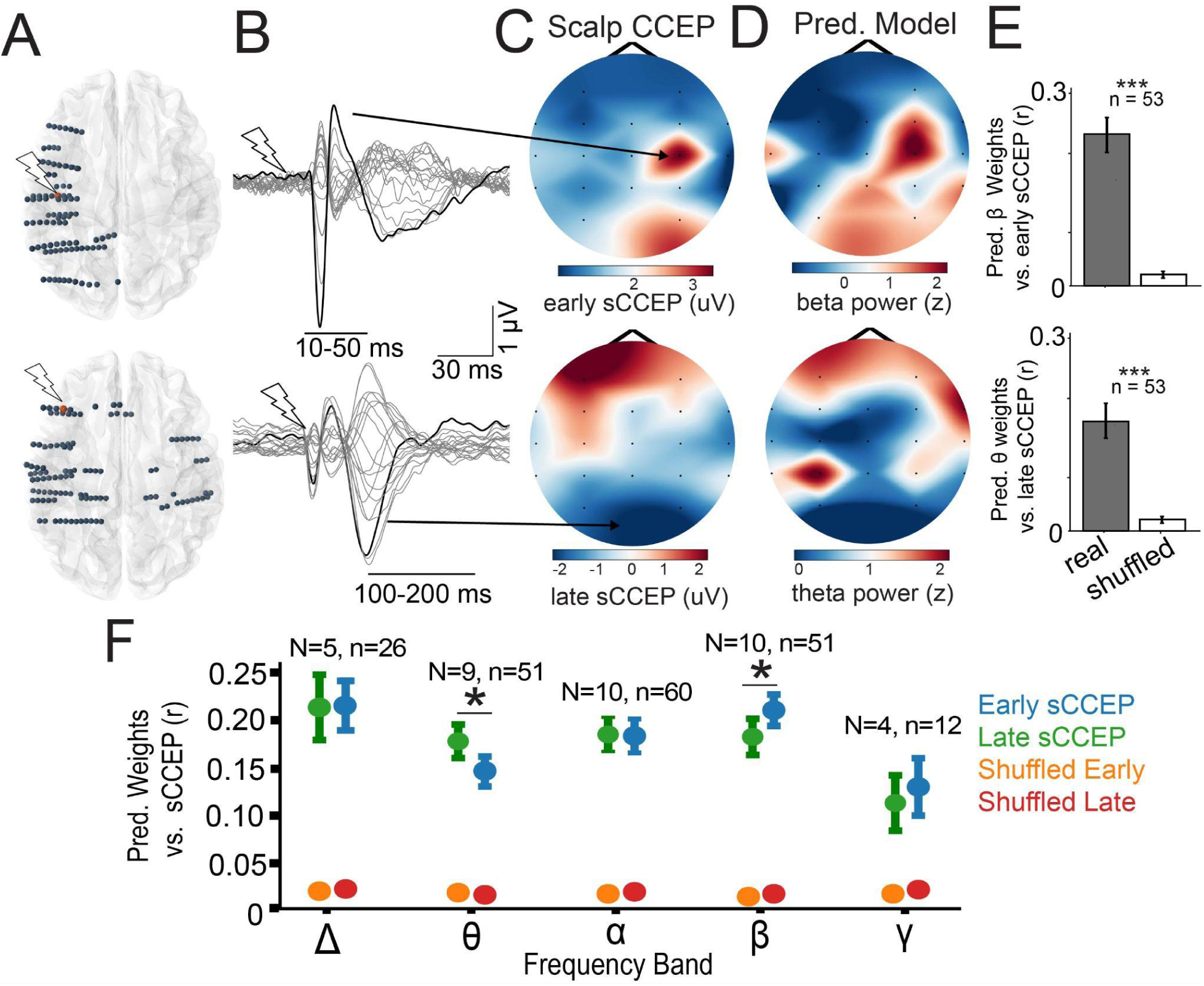
Direct electrical stimulation correlates with the predictive mapping between spontaneous scalp and intracranial activity. A) Visualization of two example stim contacts in two patients (one temporal and one frontal) B) Examples of cortico-cortical evoked potentials waveforms recorded from scalp electrodes (sCCEPs) showing early (10-50 ms) and late (100-200 ms) components at the different stimulation sites C) topomap of the early sCCEP from the top electrode in A) (top); topopmap of the late sCCEP from the bottom electrode in A) (bottom) D) Topomap showing intracranial beta power (top) and theta power (bottom) prediction weights distribution corresponding to the intracranial electrode of that row E) Comparison between the correlation of spontaneous prediction weights in beta (top) and theta (bottom) and early and late sCCEP response maps on average across all stimulation sites in all patients versus random permutation. Both correlations were significant above chance (***p<0.001, n=53 predictable contacts). F) Frequency-specific correlation coefficients between predictive maps and CCEP components in predictable contacts, with error bars indicating standard error. Late CCEP response has as significantly higher correlation than early CCEP response to theta power prediction weights (*p<0.05) and the early CCEP response has a significantly higher correlation than late CCEP response to beta power prediction weights (*p<0.05)

## DISCUSSION

### Summary of findings

In this study, we investigated whether and how intracranial brain activity can be predicted from simultaneous scalp EEG recordings in 20 patients with drug-resistant epilepsy. We hypothesized that scalp signals would best predict low-frequency intracranial activity from shallow sources, with prediction accuracy decreasing for higher frequencies and deeper sources. Our analysis revealed several key findings: 1) Intracranial activity was most predictable in lower frequencies (2-12 Hz), with scalp recordings explaining on average approximately 10% of variance across all regions and patients; the upper quartile of all spectral predictability was represented by the range 0.304 to 0.763 R^2^, consisting of 432 contacts (22.5% of total contacts), 2) Scalp signals best predicted intracranial activity in the same frequency band, particularly in theta (4-8 Hz). 3) Electrode contact depth significantly impacted predictability, with stronger predictions for shallower contacts, especially in higher frequency bands, 4) Specific brain regions showed consistent predictability across frequency bands, including temporal, parietal, and occipital areas, 5) Our leave-one-out cross-validation demonstrated that predictions generalized across patients, explaining 10-12% of variance in prefrontal, mesial temporal, and orbitofrontal regions, and 6) The validity of our predictive models was supported by correlations between model-derived scalp weights and actual responses evoked by direct electrical stimulation of intracranial sites.

Our study uniquely builds off of two key studies that also investigated simultaneous scalp and intracranial EEG. Ball et al. focused on within-patient signal-to-noise differences between SEEG and scalp EEG, while Fahimi Hnazaee et al. focused on comparing source localization derived estimates activity to intracranial activity within each patient and found weak but significant correlation in all frequency bands^27,28^. We advance this work through a substantially larger patient cohort (N = 20) with greater electrode coverage (averaging 96 contacts per patient compared to Fahimi Hnazaee et al.’s 36 contacts, which were primarily limited to hippocampal sites). Additionally, we developed computational approaches that directly predict activity across various frequency bands and brain regions, enabling assessment of both within-patient and between-patient generalizability.

### Clinical need for accurate non-invasive estimation of deep brain activity

Localizing brain activity from non-invasive scalp EEG recordings holds profound implications for understanding brain health, cognition, and behavior. Source localization technologies are particularly important for conditions where current diagnostic and monitoring tools rely heavily on subjective assessments or invasive procedures^35–37^. In neurology, for instance, epilepsy monitoring often depends on continuous scalp EEG recordings, and less commonly the use of invasive, intracranial probes for better localization of pathological activity^38,39^. In cognitive neuroscience, understanding processes such as attention or memory formation requires precise temporal and spatial resolution of neural activity. In psychiatric disorders including depression, clinical diagnosis and treatment monitoring rely primarily on expert assessments of patients’ subjective experiences, mood states, and behavioral patterns. Recent studies have shown promise in leveraging intracranial EEG signals to objectively measure these same subjective states relevant for psychiatry, such as mood fluctuations or potential for relapse in depression^40,41^. This emerging ability to capture neural correlates of the very experiences that clinicians assess demonstrates how objective neural markers could enhance clinical practice by providing complementary biological insights into psychiatric conditions. Our findings advance this goal by showing that features of intracranial brain activity in mood-relevant brain regions can be predicted from the scalp both within and between participants, underscoring the potential of noninvasive EEG recordings to infer intracranial neural dynamics. Accessible understanding of these dynamics offers potential for accelerated neuroscientific and cognitive study, as well as a pathway towards realistic personalized treatment and brain health monitoring.

### Frequency-dependent relationships and volume conduction

The variance in the predictability of intracranial activity from the scalp is closely tied to the frequency of both the iEEG and scalp EEG used as predictors (Figure 1). These nuances in predictability are represented by the matrices of R^2^ values that systematically evaluate the scalp band power input – iEEG band power output relationship. We find that intracranial activity in low frequencies, particularly in theta (4-8 Hz), are most predictable (Figure 1 G-I). We also find that the scalp signals that are most predictive of intracranial signals are those recorded at the same frequency (Figure 1G,J right). These trends may be explained by characteristics of neural signal propagation and transmission that differ across frequency bands. For example, in volume conduction models of brain activity, high-frequency signals are more localized, hence harder to predict from surface recordings, while low-frequency signals, capable of traversing larger brain volumes, are more easily detected on the scalp^30,31^. Thus, it may follow that a linear combination of similar frequency signals are best suited to reconstruct an original deeper laying signal, explaining the relative predictive success of like-band regressions. However, at the same time, our analysis across these various frequency bands suggests unique roles and propagation characteristics for each. Delta (2-4Hz) and theta (4-8 Hz) frequencies, showing higher predictability, may reflect the notion that more global neural processes or synchronized activity across larger brain networks may be mediated through these frequencies^32,42^. Alpha (8-12 Hz) band predictability might indicate the involvement of cortical idle or inhibitory processes with modest regional specificity^43,44^. Beta (12-30 Hz) frequencies, with their lower predictability, and gamma frequencies (30-48 Hz) with near zero predictability, likely mirror more localized or function-specific brain activities.

### Spatial and depth considerations of scalp-to-intracranial prediction

At the same time, the predictability of activity from specific intracranial contacts is significantly influenced by spatial factors, such as the contact’s depth and its location within the brain. Our analysis revealed consistent regional patterns of predictability across multiple frequency bands, with particularly strong average predictions in temporal (R² ranging from 0.15 - 0.30), parietal (R² ranging from 0.17 - 0.35), frontal areas (R² ranging from 0.28 - 0.32). The insula and middle temporal regions showed notably high predictability in lower frequency bands (delta and theta), while parietal regions demonstrated stronger predictability in comparatively higher frequencies (alpha and beta) (Figure 2). In addition, we find that in all frequency bands, predictability decreases as a function of the contact’s depth, defined as its distance away from the nearest scalp electrode. This effect is most pronounced in the beta band across all patients and all contacts and in the delta, alpha, and beta bands across all contacts in the frontal lobe. The depth-dependent effect was particularly evident in alpha and beta bands, where predictable contacts were significantly shallower than unpredictable ones (differences of 1.9mm and 4.5mm, respectively). Notably, while theta activity is most predictable across all contacts in all patients, it also shows no significant increase or decrease in predictability as a function of contact distance. This may signal that theta activity predictability stems from global oscillatory processes that are preserved across varying depths of brain^45–47^, unlike other frequency bands, which show more depth-dependent attenuation characteristics.

### Hierarchical Bayesian analysis of predictability factors

Next, our hierarchical Bayesian analysis affords a more nuanced understanding of predictability, explaining to the best of our ability the effect of depth, frequency, absolute power, and relative strength of the signal (normalized band power), establishing posterior beliefs on the predictability of a new contact in a new patient. First, we confirm the trends made in the more traditional frequentist analysis above about the effect of depth on predictability, indeed seeing a negative coefficient associated with increased depth across all predictable frequency bands (Figure 3E). Next, the negative effect between absolute power and predictability across all frequencies, suggests that contacts capturing high overall neural activity might also pick up more noise or non-specific activity, diluting the predictability of our like-band regression models that predict specific frequency bands. (Figure 3F) Conversely, the positive correlation of normalized relevant frequency power with predictability indicates that contacts that are sampling sources that are strong and specific oscillators tend to be more predictable. In contrast, contacts that are sampling a diversity of sources and oscillators tend to be less predictable in any one given frequency band (Figure 3G). This makes intuitive sense, and may warrant further feature selection for certain contacts where this particular spectral decomposition of the signal may not be providing the most information for linear prediction. Of note, both of these effects are more strongly pronounced in the prediction of theta. This could be for two reasons: 1) our model is only as good as our data, and due to theta oscillations being most predictable and having more variance across all patients, we can make stronger conclusions about these effects and 2) theta activity is particularly susceptible to both specific and non-specific activity within its spectral window. This could explain why a decrease in absolute band power, signaling potentially more non-specific activity, leads to disproportionately less predictability, relative to other bands, while an increase in normalized band power within theta corresponds to disproportionately more predictability. We extend our major takeaway that there may be evidence suggesting certain generalizable factors in modeling intracranial predictability by explicitly testing it via Leave-One-Out cross validation. These models demonstrate that intracranial activity predictability is generalizable across patients for regions of interest (ROIs). This suggests the existence of common pathways for signal propagation from invasive to non-invasive recordings across patients, with the most successful inference occurring in the prediction of theta activity (Figure 4B, D, F). The prominence of scalp electrodes closest to an RoI in predicting activity further underscores the spatial dependency of our models, though with variations across different frequencies (Figure 4C, E, G).

### Study limitations and future directions

While our study provides foundational insights into the predictability of intracranial activity from scalp recordings, several limitations warrant mention. First is the generalizability of our findings across broader populations and conditions. Our study was limited by the presence of epilepsy in our patients. We were rigorous in our treatment both in the preprocessing and analysis to limit the influence of pathological signals by removing pathological contacts and contacts identified independently by neurologists. As a final check, we also visually inspected the band-passed theta signal noting, where studies have shown relationships between IEDs and abnormal theta waveform or power^48,49^. After preprocessing, we noted no strong outliers or obvious epileptic activity in each patient upon visual inspection of each contacts’ spectra. Additionally, we compare the trends in predictability across the subset of the participant population with MRI-positive epilepsy and find no significant differences between that subset and the MRI-negative subset (Figure S3). Nevertheless, future studies leveraging more advanced pathological signal removal techniques or non-epilepsy patient populations with intracranial contacts^36,50^ would provide insight as to the generalizability of these findings. Furthermore, the relationship between scalp and intracranial recordings may be altered in all patients due to surgical changes to the skull, including the implanted contacts and drilled holes, potentially limiting generalizability to nonsurgical patients. Next, our study is limited by the specificity of predictions to certain brain regions due to heterogeneous contact placement. Future studies with larger contact coverage per patient or with more sophisticated methodologies for extrapolating accurate source locations with sparse sampling may improve predictability and spatial resolution and specificity of those predictions. Finally, there is the potential for refining our statistical models with more sophisticated algorithms or additional variables. Although the observed average explained variance (R^2^ between 0.1-0.2, top quartile R^2^ between 0.3-0.73) represents a level of predictability higher than historical attempts, which are on average approximately ∼R^2^ = 0.05 ^27–29^ to recover intracranial brain activity from scalp sensing, seeking stronger relationships could be relevant for potential applications of reconstructed intracranial signals. Of note, high frequency activity (gamma and high gamma) have been studied in depth as a promising signal to relate to neuronal activity^51–53;^ thus advances in improving the prediction of that activity, which we showed was mostly unpredictable within this study’s methods, is also needed. As an augmentation, the exploration of non-linear models or deep learning approaches could offer deeper insights into brain activity propagation, though these approaches require larger datasets to capture neural patterns effectively. Due to their numerous parameters, these models require a substantial number of examples to learn patterns while avoiding overfitting.

In conclusion, our study advances the understanding of the relationship between noninvasive EEG signals and deeper brain activities with both extensive coverage and larger sample size, focusing on the spatially- and frequency-dependent nature of this relationship. By constraining our modeling to linear relationships, we provide the most interpretable investigation of simultaneous invasive and non-invasive neural activity to date. We intend for these findings to contribute to the development of objective, noninvasive biomarkers for psychiatric and neurological conditions and lay the foundation for future studies in this topic area.

## METHODS

### Participants

20 patients (median age = 32 ± 8.03, 10 female, Table S1) from the Claudio Munari Epilepsy Surgery Center of ASST GOM Niguarda Hospital, (Milan, Italy) participated in this study. All participants had a history of drug-resistant focal epilepsy, and they underwent stereo-electroencephalography (SEEG) contact implantation for seizure localization and possible surgical removal/ablation of the seizure onset zone (SOZ). As determined by a neurologist, 14 participants did not show any anatomical malformation/lesion on MRI, while 6 showed small anatomical alterations (see Table S1 for description of pathology, etiology, and clinical assessments). The location of contact placements were decided purely based on clinical reasons. All participants provided their Informed Consent in accordance with the local Ethical Committee (ID 348-24.06.2020, Milano AREA C Niguarda Hospital, Milan, Italy) and with the Declaration of Helsinki.

### Electrode placement and localization

Details for SEEG contact placement and localization are reported in previous studies^34,54,55^. In summary, for each participant, preoperative MRI (with Achieva 1.5 T, [Philips Healthcare]) and CT (O-arm 1000 system [Medtronic]) were obtained for SEEG planning. SEEG shafts were placed by the surgeon using a robotic assistant (Neuromate, Renishaw Mayfield SA) under general anesthesia. SEEG contacts consisted of a variable number (from 13 to 19, mean=16.4) of platinum–iridium semi flexible multi-contact intracerebral contacts, with a diameter of 0.8 mm, a contact length of 2 mm, an inter-contact distance of 1.5 mm and a maximum of 18 contacts per contact (Microdeep intracerebral contacts, D08 [Dixi Medical]). After implantation, a fine cone-beam CT data set was acquired by using the O-arm and coregistered with the T1-weighted 3D MR using FLIRT^56^ . Electrode positions and anatomical labels were obtained using Freesurfer (Desikan-Killiany Atlas)^57–59^ and SEEG-Assistant^60^. Normalized coordinates were obtained by performing a non-linear registration between the participant’s skull-stripped MRI and the skull-stripped MNI152 template (ICBM 2009a Nonlinear Symmetric) using ANTs’ SyN algorithm^57,61^.

### Simultaneous SEEG and hd-EEG recordings

SEEG activity from implanted patients was continuously recorded through a 192-channel recording system (Nihon-Kohden NeuroFax-1200) with a sampling rate of 1000Hz. All acquisitions were referenced to two adjacent electrode contacts located entirely in white matter. During their last day of hospitalization, all participants in the present study underwent simultaneous scalp non-invasive recordings through high-density 256 channel electroencephalography (hd-EEG Geodesic Sensor Net, HydroCel CleanLeads) as in^34,55^ Parmigiani et al., *Brain Stimulation*, 2022; and Mikulan et al., *Scientific Data*, 2020. Placement of the hd-EEG net on the head was performed by trained neurosurgeons using sterile technique, following a precise step-by-step protocol: (1) sterilization of the net, (2) removal of the protective bandage from the participant’s head, (3) skin disinfection with Betadine and Clorexan, (4) positioning of the hd-EEG net, and (5) reduction of the impedances below 25–50 kOhm using conductive gel. An example of this setup is shown in Fig. 1a. Hd-EEG was then recorded at a 1000 Hz sampling rate using an EGI NA-400 amplifier (Electrical Geodesics, Inc; Oregon, USA) referenced to Cz. SEEG and hd-EEG recordings were aligned using a digital trigger signal generated by an external trigger box (EMS s.r.l., Bologna, Italy). At the end of the simultaneous data acquisition, the spatial locations of hd-EEG electrodes and anatomical fiducials were digitized with a SofTaxicOptic system (EMS s.r.l., Bologna, Italy) and coregistered with a pre-implant 3D-T1 MRI. The net was then removed, and the skin was disinfected again.

### Data pre-processing

For both SEEG and scalp EEG, preprocessing included bad channel removal, notch filtering, detrending, demeaning, and referencing. As underlying muscle and eye movements influence scalp EEG, an additional independent component analysis (ICA) step was applied to remove those components. The specifics for each preprocessing procedure are outlined below (and in Supplementary Figure 1).

#### SEEG data

Before preprocessing, SEEG contacts in or near the epileptic focus as determined by the clinical team for each participant were removed from the analysis. SEEG recordings were then bipolar rereferenced as is relatively standard in SEEG recordings^62,63^. To do so, we calculated the difference between adjacent contact pairs along each depth contact to minimize common electrical noise and maximize spatial specificity of each signal. These bipolar electrode contact pairs are referred to as “contacts” throughout analyses here for brevity. Data were then demeaned (the mean of each signal is subtracted, yielding a zero-mean for each contact) and detrended (1 Hz high pass filter, 3rd order Butterworth) to eliminate low frequency drift. Next, a 50 Hz notch filter was applied to eliminate line noise. SEEG recordings were inspected to reject contacts that contained variance greater than one standard deviation above the mean variance of all contacts in a patient. Continuous recordings were then bandpass filtered using MATLAB’s ‘bandpass’ function with default parameters (minimum-order Butterworth filter with 60 dB stopband attenuation) into canonical bands of interest into canonical bands of interest (delta: 2-4 Hz, theta: 4-8 Hz, alpha: 8-12 Hz, beta: 12-30 Hz, gamma: 30-48 Hz, high gamma: 52 - 100 Hz)). We limited the delta band to 2 hz to avoid overlap with the detrending preprocessing step which high passes the signal at 1 Hz. For gamma and high gamma bands, 48 to 52 Hz were avoided to ensure no overlap with the filtered out 50 Hz line noise. Bandpassed recordings were then split into 1 second, non-overlapping epochs which were selected as a balance of containing both low- and high-frequency information in a single epoch. Data were visually inspected for channels with signs of electrical artifact or beyond threshold noise, but no channels were deemed necessary for removal.

#### Scalp EEG

First, bad channels were automatically identified via a thresholded data-driven Wiener estimation using the TESA toolbox^64,65^. Bad channels were then interpolated via spherical interpolation using EEGLAB^66,67^. Data were then detrended (1 Hz high pass filter, 4th order Butterworth filter). Next, we performed independent component analysis (ICA) and automatically labeled components for rejection using ICLabel^68^ to remove non-neural signals such as eye blinks, movements, and muscle activations. Next, data were rereferenced to maximize spatial specificity of each individual sensor and spatially downsampled in order to limit the dimensionality of input features and minimize common noise across scalp electrodes. To do so, at each of 19 target sensors corresponding to locations on the 10-20 contact map (for maximal translation), a local spatial signal was obtained by subtracting the average signal of the five nearest neighbor sensors from the target sensor signal. This laplacian rereferencing scheme^66,69^ was chosen in order to both leverage the large coverage of the 256ch EEG cap while yielding a set of features that is both relatable to out-of-study data sets using less advanced headsets and containing an input feature size appropriate for out amount of data. This procedure left 19 rereferenced scalp channels for each participant. Continuous recordings were then 50 Hz notch filtered and demeaned with the same technique as the SEEG data. Continuous recordings were then bandpass filtered (identical parameters as above) into canonical bands of interest^70^ (delta: 2-4 Hz, theta: 4-8 Hz, alpha: 8-12 Hz, beta: 12-30 Hz, gamma: 30-48 Hz). Bandpassed recordings were divided into non-overlapping 1-second epochs, with scalp EEG and SEEG epochs temporally aligned. Each scalp EEG epoch was then examined for electrical artifacts. Epochs were rejected if any channel exceeded a predefined amplitude threshold indicating non-neural activity (5.3 ± 2 (∼1%) epochs removed across participants). The corresponding time-matched SEEG epochs were also removed when artifacts were detected in the scalp recordings. Notably, while scalp EEG epochs were screened for artifacts, no additional artifact rejection was performed on the SEEG recordings since no clear artifacts were visible upon visual inspection. The cleaned SEEG and scalp EEG datasets were then used for feature extraction and statistical modeling.

### Feature extraction

For each SEEG and scalp EEG time series, average bandpower for each 1 second non-overlapping epoch was calculated at each frequency band of interest by summing the square amplitude of the band-passed time series at each epoch using the bandpower function in MATLAB. These continuous band power time series, where each value represents the band power calculated in that 1 second epoch, are then converted to log-power via natural logarithm in order to increase normality. The log-power time series are then Z-scored to baseline mean log-power at the single contact level. These Z-score time series were then used as the input (scalp EEG) and output (intracranial SEEG) features for statistical modeling. Of note, when analyzing models using features captured from a shorter time window (500 ms), we also observed similar results with respect to the spectral predictability trends observed with 1 second epochs (Fig S4). For spatial labeling, recording sites were organized into regions and parcellations according to the Desikan-Killiany atlas^58^ and the location of their centers in MNI space^61^. The three MNI-centered regions of interest we focused on here were the mesial temporal lobe (defined as a 30 mm radius around the amygdala MNI coordinate (-27, -4, -20), the orbitofrontal cortex (defined as a 30 mm radius around the subcallosal cingulate cortex MNI coordinate (-6, 24, -11) and the prefrontal cortex (defined as a 30 mm radius around the left dorsolateral prefrontal cortex MNI coordinate (-30, 43, 23). The 30 mm radius was chosen as a balance between enough sampling contacts across patients and focality of the defined region.

### Statistical Modelling

The present study used multiple statistical models to investigate the predictability of intracranial neural activity compared to scalp neural activity. They are presented in order of appearance.

#### Regularized Linear Regression

First, this study utilized ridge regression^71^, a common technique to limit overfitting of linear regression, the predictive statistical model. For each participant, scalp EEG the power time series in each frequency band (19 contacts x 1 frequency x t seconds) are regressed against each SEEG contacts band power time series (1 SEEG contact x 1 frequency x t seconds). This yields a set of i*j regressions where i represents the number of input bands analyzed and j represents the number of output bands analyzed (in this case, i = j = 5) per SEEG contact per patient. This approach was used to easily interpret frequency specific predictability at each intracranial contact. For each outer cross-validation fold, the data is split into a training set consisting of a contiguous stream of the original data about 80% of the length of the original data. The test set is then taken as the remaining 20% of the data with a 6 second lag removed on each side of the training set. This methodology ensured that the training data set does not implicitly share data with the test set via autocorrelation, which could be present if the epochs selected for training and test were selected randomly. This time window (6s) was chosen via analysis of the average partial autocorrelation function. An additional inner 10-fold cross validation for each selected 80-20 split was used for hyperparameter tuning of the regularization constant. The outer 10-fold validation varying each 80% training set was used to generate a confidence interval for R^2^ values of each individual regression. The ridge regression model was fitted using the python package Sci-kit learn. Ridge regression was chosen over alternatives such as elastic net and Lasso due to improved computation costs and a lack of a need for sparsity associated with such methods.

#### Hierarchical Bayesian Linear Regression

In order to assess the effect of a variety of potentially confounding parameters on the predictability of a given intracranial contact, we utilized a Hierarchical Bayesian linear regression^33^. This approach was chosen for multiple reasons. Firstly, this method allows us to take into account that our data involves repeated samples from a limited group of individuals. By structuring the model hierarchically, we can make inferences on global population parameters, representing the effect that variables have on predictability in new individuals not included in the studies. This is critical, as our ultimate scientific goal is to make statements about how these parameters may generalize *to new patients*. Bayesian inference naturally allows information to “flow” from the individual-level observations to the population parameters. Second, the incorporation of prior information in the form of prior distributions on the parameters reduces variance in the parameter estimates. This is particularly critical due to the previously-mentioned limited clinical sample size. The lower-level parameters in this setup correspond to individual-specific effects, while higher-level parameters correspond to the population effects of interest. The key parameters (contact depth, normalized spectral power of the predicting band, and absolute broadband power of the contact) were selected to hone in on three key hypotheses. First, as introduced earlier, there seems to be a strong depth dependence on predictability, especially in higher frequencies; however, due to variation in contact region and the heterogeneous distribution of contacts, it is imperative to better characterize how important depth is to the prediction of the contact. Second, the normalized spectral power of the predicting band represents a functional signal-to-noise ratio for that prediction. High normalized power in a given contact signifies strong sampling of a source that is actually oscillating at the given frequency band, while low normalized power could signify strong noise from off-frequency sources. Finally, absolute broadband power gives us a measure for non-frequency specific sources of signal or noise, and its inclusion as a model parameter is to disambiguate those sources’ effect on predictability. In other words, predictability of any given frequency band being strongly related to the absolute broadband power of the contact might indicate that linear models are picking up off-target or non-generalizable information such as pathological events or recording artifacts. The model infers each of these parameters as a distribution for each regression coefficient. The posterior distributions for each patient describing each of the regression coefficients and intercept is presented in Figure S6.

Our probabilistic model is defined as μ_α_ ∼ N(0, 100.0)

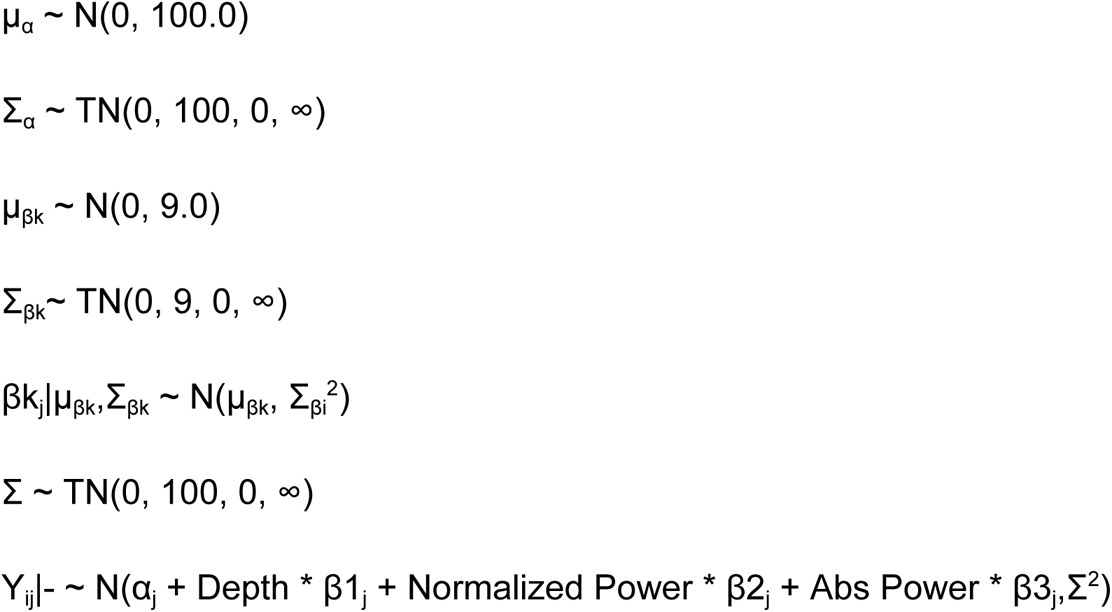

(the notation *|- means conditioned on the remaining variables). Here, Y refers to the inferred R^2^ of a ridge regression of a pair of input and output features of a contact (for example, the R^2^ reflecting the predictability of a given contact’s theta power as predicted by scalp theta power). Our parameters of interest are the “global means” (μ_β1_, μ_β2_, μ_β3_), which represent the population (new person in clinic) predictability in terms of the specific covariate. Returning to the primary motivation for using a hierarchical model, a large number of samples in a single individual j will result in precise inferences on α_j_, β1_j_j, β2_j_, and β3 _j_but with limited influence on our global parameters of scientific interest.

Inference was performed using the Numpyro package in python. This package uses Hamiltonian Monte Carlo to draw samples from the posterior distribution (Betancourt 2017). Functions evaluated on the samples (such as quantiles) from the posterior can approximate the same function on the posterior. We used 2000 samples with an additional 500 burnin samples. We used the No U-Turn Sampler (NUTS) with an acceptance probability of 0.8 and step size of 1.0 corresponding to default values.

#### Leave-one-out Ridge Regression

Finally, in order to gauge the generalizability of predictions of intracranial activity from the scalp, we utilized a Leave-one-patient-out cross validation scheme with ridge regression. In these models, first regions of interest were selected and the subset of intracranial contacts from each patient that fulfilled a distance criteria were used as the data for each model. Specifically, the mesial temporal lobe (defined as a 30 mm radius around the amygdala MNI coordinate (-27, -4, -20), the orbitofrontal cortex (defined as a 30 mm radius around the subcallosal cingulate cortex MNI coordinate (-6, 24, -11) and the prefrontal cortex (defined as a 30 mm radius around the left dorsolateral prefrontal cortex MNI coordinate (-30, 43, 23) were selected. Then, for each region, the synchronized simultaneous scalp and intracranial data are then used as the total dataset for each model. The input and output space was similarly implemented to the ridge regression model above, where data from a single scalp band from all 19 scalp electrodes are regressed against a single intracranial band. However, rather than looking at an individual patient, the training set is formed from all epochs in n-1 patients, and the test set is all epochs of a left out patient (which may include a number of different contacts contained within the region of interest). This process was then repeated for each patient. In other words, we construct models that predict a left-out patient’s intracranial contact activity from their scalp activity at each pair of spectral features. We then average the results of all runs, making each presented result the average of n models, each tested on a different left-out patient, where n is the number of patients with contacts implanted in the specified region of interest. Within each Leave-One-Out loop, an identical method for hyperparameter tuning as above was used. The final evaluation of each model was reported as the average of R2 values and average of regressions coefficients across all patients. The ridge regression model was done using the python package Sci-kit learn.

### Statistical Analysis

Differences in mean model performance within participants between groups of contacts as well as differences in mean model performance between participants at the group level were assessed with the non-parametric Mann-Whitney U Test. Our specific hypotheses for these statistical tests are elaborated in Fig 1. Determination of significantly predictable contacts, as presented in Figures 1 and 2, was done by calculating a 95% confidence interval of R^2^ values through 10-fold cross validation in each within-patient Ridge Regression (see above *Regularized Linear Regression*). For a given regression, intracranial contacts with mean R^2^ greater than 0 and with the lower bound of the 95% confidence interval greater than 0 were classified as “predictable”, and contacts failing that criteria were deemed as “not predictable”. Correlation coefficients between intracranial contact distance and predictability, as well as scalp electrode distance and regression coefficient (weight) were calculated using least squares linear regression. The average R^2^ of predictable contacts across each same band-same band regression at each spatial parcellation was then mapped (Figure 2A, tabulated further in Supplementary Table 2). Significance of group mean correlation coefficients with distance were assessed using one-sample t-tests. Likewise, in the Leave-One-Patient-Out cross-validation study, significance of each mean R2 value was tested with 1-sample t-tests, and significance of relationship of model regression coefficients to scalp-to-intracranial euclidean distance were calculated using simple linear regression with scipy linregress function (significance is a two-tailed T-test with null hypothesis being the slope is equal to 0). All statistical analysis was performed using python packages scipy and sci-kit learn^72,73^. All contact and head visualization was performed through MNE-python^74^.

## Acknowledgements

The authors are grateful for the participants that dedicated their time to this research. We extend our deepest gratitude to the patients who participated in this study, as well as the nursing and physician staff at Niguarda Hospital in Milan, Italy. We would also like to give special thanks to Manish Saggar, Laura Gwilliams, Guosong Hong, Alberto Salleo, Umair Hassan, Scott Linderman, and Will Giardino for their valuable feedback. Special thanks also to Jade Truong for her support throughout the research process. Formal credit: **Ajay K. Subramanian**: Conceptualization, Methodology, Software, Validation, formal data analysis, Writing – review & editing. **Austin Talbot**: Conceptualization, Methodology, Writing – review & editing, Software **Naryeong Kim**: Data curation, data analysis. **Sara Parmagiani**: Methodology, Writing review & editing, Resources, Supervision. **Christopher Cline**: Conceptualization, Methodology, Software, Writing - review & editing, Supervision **Ethan Solomon**: Writing review & editing **James Winn Hartford**: Data curation, Writing - review & editing, **Yuhao Huang**: Writing - review & editing. **Ezequiel Mikulan**: Methodology, Review, Resources. **Flavia Maria Zauli**: Data curation, review & editing. **Francesco Cardinale**: Patients’ clinical management. Piergiorgio d’Orio: Patients’ clinical management. **Francesca Mannini**: Writing - review & editing**. Andrea Pigorini**: Methodology, Writing – review & editing, Resources, Supervision. **Corey J. Keller**: Conceptualization, Methodology, Writing – review & editing, Resources, Data curation, Supervision.

