## Supplemental Figures and Tables for "Scalp EEG predicts intracranial brain activity in humans"

### Table S1: Pathology and Clinical information for the patient population

| Patient | Sex | Age | MRI | Side of SEEG | Lobes of SEEG | Therapy | N° electrodes |
|---|---|---|---|---|---|---|---|
| **1** | M | 19 | Negative | Bilat | Right inferior frontal gyrus + Left fronto-central lobes | CBZ 1200 mg/die | 14 (12 left + 2 right) |
| **2** | M | 46 | Negative | Bilat | Bilateral central and temporo-perisylvian lobes | CBZ 800 mg/die; LRZ 1mg/die; LEV 2000mg/die; Risperidone 4 mg/die | 18 (13 right + 5 left) |
| **3** | M | 19 | Negative | Right | Right temporal and parieto-occipital lobes | CBZ 1200 mg/die; LEV 3000 mg/die; PGB 75 mg/die | 17 |
| **4** | F | 20 | Negative | Left | Left temporo-parieto-perisylvian lobes and inferior frontal gyrus | LTG 200 mg/die; CLB 10 mg/die | 14 |
| **5** | F | 28 | Negative | Bilat | Right temporo-perisylvian lobes and inferior frontal gyrus + Left temporo-fronto-central and perisylvian lobes | LCM 400 mg/die; OXC 600 mg/die | 16 (10 left + 6 right) |
| **6** | M | 36 | Negative | Left | Left temporo-perisylvian lobes | CBZ 1400 mg/die; LEV 3000; LCM 400 mg/die | 14 |
| **7** | F | 40 | Negative | Left | Left temporo-parieto-occipital lobes | LTG 400 mg/die; TPM 100 mg/die | 16 |
| **8** | M | 33 | Bilateral PNH | Bilat | Right temporo-perisylvian and occipital lobes + Left temporo-occipital lobes | CBZ 800 mg/die | 18 (15 right + 3 left) |
| **9** | F | 34 | Right PNH | Right | Right temporo-occipito-parietal lobes | CBZ 1000 mg/die; LEV 2500 mg/die | 13 |

| | | | | | | | |
|---|---|---|---|---|---|---|---|
| **10** | F | 46 | Left temporo-basal FCD | Left | Left temporo-occpital lobes | LCM 400 mg/die | 17 |
| **11** | M | 46 | Bilateral PNH | Bilat | Right temporo-occipital lobes + Left temporal lobe | CBZ 1400 mg/die; PB 100 mg/die | 17 (11 right + 6 left) |
| **12** | M | 35 | Negative | Right | Right fronto-centro-temporal lobes | CBZ 1400 mg/die; LEV 750 mg/die | 19 |
| **13** | F | 31 | Negative | Bilat | Right temporo-perisylvian lobes + Left temporal lobe and inferior frontal gyrus | CBZ 1200 mg/die; TPM 150 mg/die | 15 (10 right + 5 left) |
| **14** | M | 32 | Negative | Right | Right fronto-centro-temporal lobes | LEV 3000 mg; LCM 300 mg | 15 |
| **15** | F | 31 | Negative | Right | Right temporo-parieto-occipital lobes and inferior frontal gyrus | LEV 2000 mg/die | 15 |
| **16** | M | 23 | Negative | Left | Left temporo-perisylvian and inferior frontal gyrus | ZNS 200 mg/die; PB 150 mg/die | 13 |
| **17** | F | 29 | Left fronto-insular atrophy + Left HS | Left | Left fronto-centro-parieto-perisylvian and temporal lobes | CBZ 1200 mg/die, PER 6 mg/die | 15 |
| **18** | M | 44 | Negative | Right | Right temporo-parieto-occipital lobes and superior frontal lobe | CBZ 1400 mg/die; PB 100 mg/die; LTG 400 mg/die | 17 |
| **19** | F | 27 | Left superior temporal FCD | Bilat | Left temporo-perisylvian lobes + Right superior temporal gyrus | LCS 350 mg/die; PER 6 mg/die; CLZ 0.5 mg/die | 17 (15 left + 2 right) |
| **20** | F | 36 | Negative | Left | Left fronto-temporo-perisylvian lobes | CBZ 1400 mg/die; LTG 400 mg/die | 14 |

Bilat: Bilateral; BRV: Brivaracetam; CBZ: Carmabazepine; CLB: Clobazam; CLZ: Clonazepam; ESL: Eslicarbazepine acetate; FCD: Focal Cortical Dysplasia; HS: hippocampal sclerosis; LCM: Lacosamide; LEV: Levetiracetam; LRZ: Lorazepam; LTG: Lamotrigine; OXC: Oxcarbazepine; PB: Phenobarbital; PER: Perampanel; PGB: Pregabalin; PNH: periventricular nodular heterotopia; TPM: Topiramate; VPA: Valproate; ZNS: Zonisamide

**Table S2: Low frequency and diagonal regressions hypothesis test table for example patient in Figure 1 and for all patients.** Values with R2 > 0.1 are highlighted

| | | Scalp Frequency Regressions | | | | Intracranial Frequency Regressions | | | | Diagonal vs Off-Diagonal | | | |
|---|---|---|---|---|---|---|---|---|---|---|---|---|---|
| | | Mean $R^2$ ± SD (n) | | p-value | Effect Size | Mean $R^2$ ± SD (n) | | p-value | Effect Size | Mean $R^2$ ± SD (n) | | p-value | Effect Size |
| Patient Group | Contact Type | Low Freq | High Freq | | | Low Freq | High Freq | | | Diagonal | Off-Diagonal | | |
| Example Patient | All Contacts | 0.095 ± 0.14 (n = 1,650) | -0.007 ± 0.09 (n = 1,100) | p < 1e-9 | 0.40 | 0.09 ± 0.15 (n = 1,650) | 0.003 ± 0.09 (n = 1,100) | p < 1e-9 | 0.28 | 0.12 ± 0.16 (n = 550) | 0.04 ± 0.12 (n = 2,200) | p < 1e-9 | 0.20 |
| | Predictable Contacts | 0.24 ± 0.11 (n = 629) | 0.19 ± 0.07 (n = 120) | p = 0.0011 | 0.12 | 0.25 ± 0.10 (n = 596) | 0.16 ± 0.07 (n = 153) | p < 1e-9 | 0.35 | 0.27 ± 0.13 (n = 222) | 0.21 ± 0.09 (n = 527) | p = 1.4e-8 | 0.20 |
| All Patients | All Contacts | 0.00 ± 0.15 (n = 28,770) | -0.04 ± 0.12 (n = 19,180) | p < 1e-9 | 0.14 | -0.01 ± 0.15 (n = 28,770) | -0.03 ± 0.12 (n = 19,180) | p < 1e-9 | 0.07 | 0.02 ± 0.16 (n = 9,590) | -0.02 ± 0.13 (n = 38,360) | p < 1e-9 | 0.11 |
| | Predictable Contacts | 0.25 ± 0.12 (n = 4,393) | 0.19 ± 0.10 (n = 1,266) | p < 1e-9 | 0.17 | 0.25 ± 0.12 (n = 4,161) | 0.19 ± 0.10 (n = 1,498) | p < 1e-9 | 0.23 | 0.27 ± 0.14 (n = 1,774) | 0.22 ± 0.10 (n = 3,885) | p < 1e-9 | 0.17 |

**Note: Higher $R^2$ values and effect sizes indicate stronger effects. Values with $R^2 > 0.1$ are highlighted.**

Statistical significance: Mann-Whitney U test p-values are shown, with $p < 0.05$ considered significant.

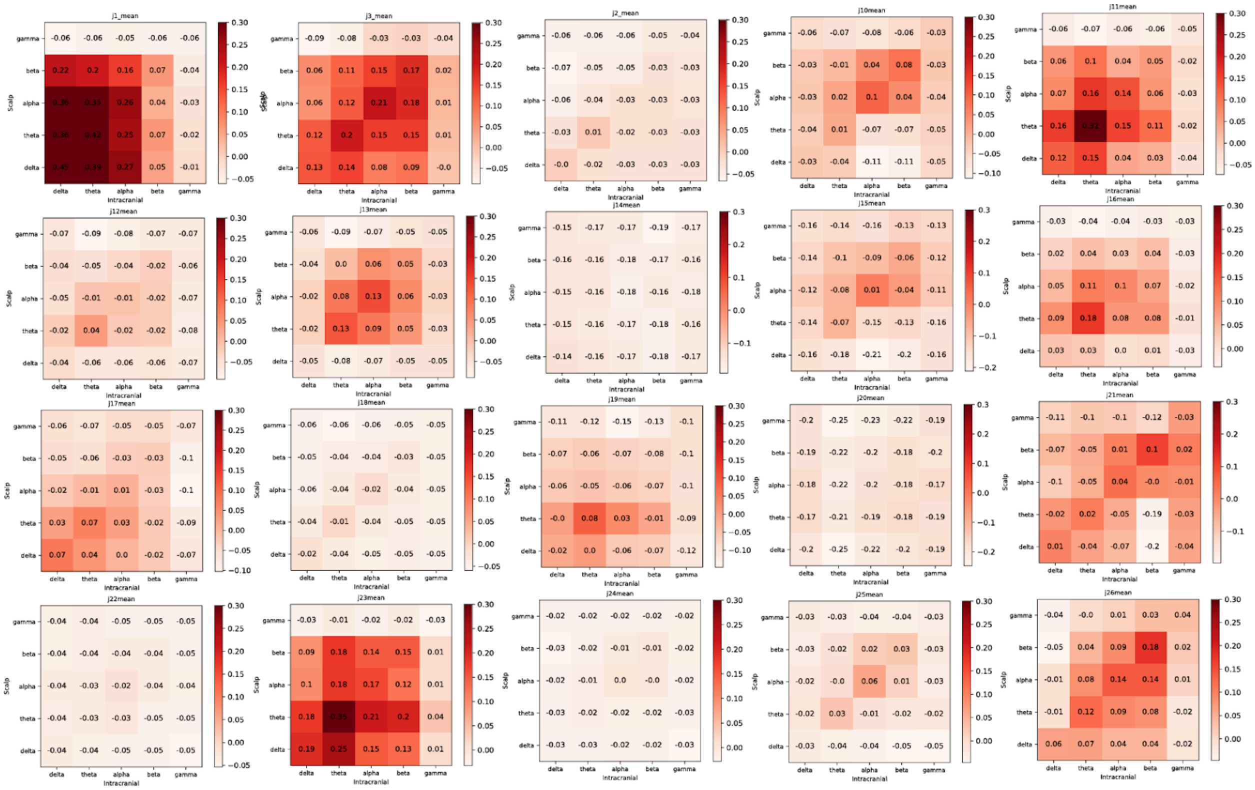

**Figure S1: Mean $R^2$ heatmaps for all contacts within each patient.**

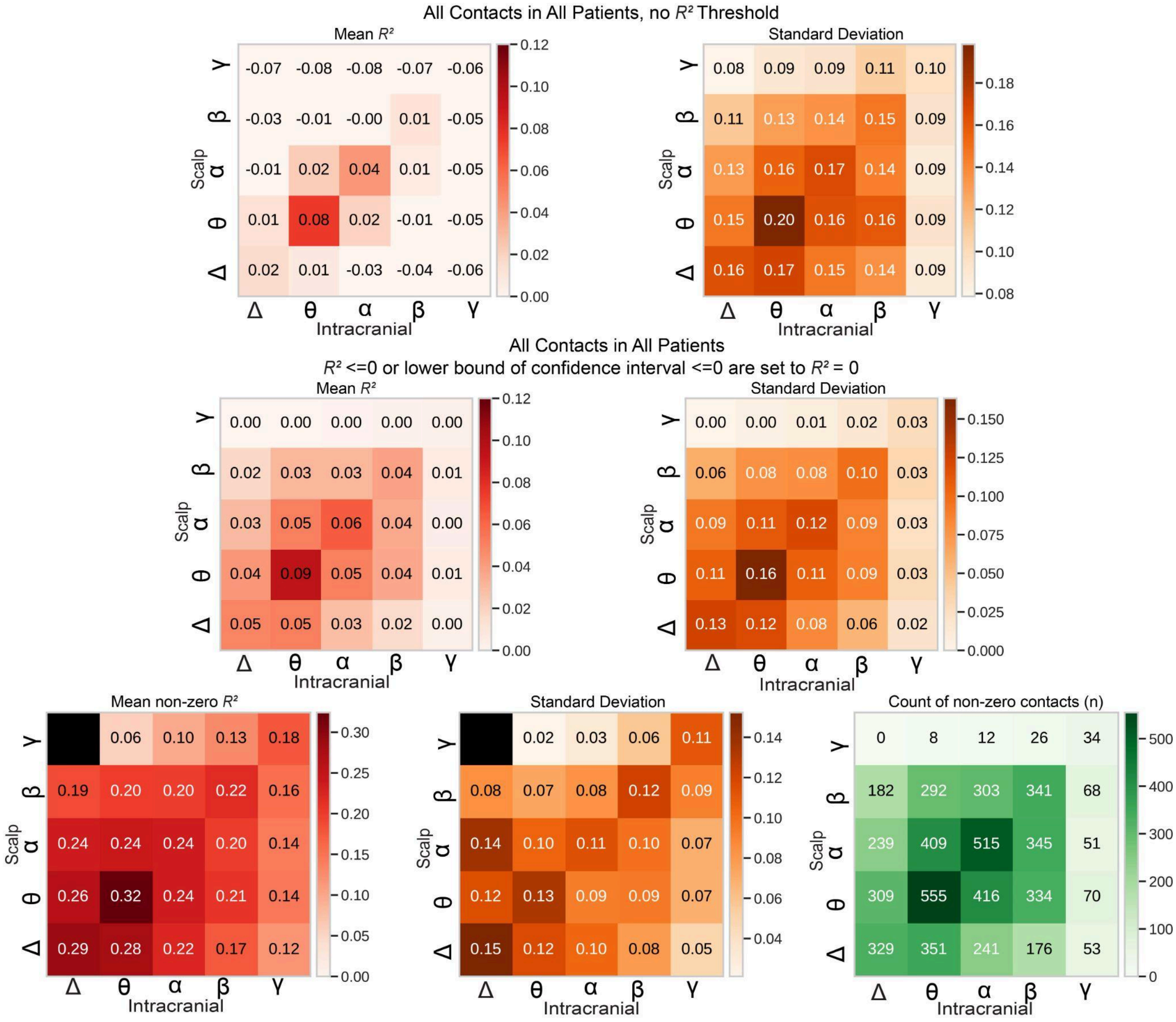


**Figure S2: Mean $R^2$ Heatmaps for all contacts in all patients with raw $R^2$ values, with unpredictable contacts set to $R^2$ = 0, and for only predictable contacts.**

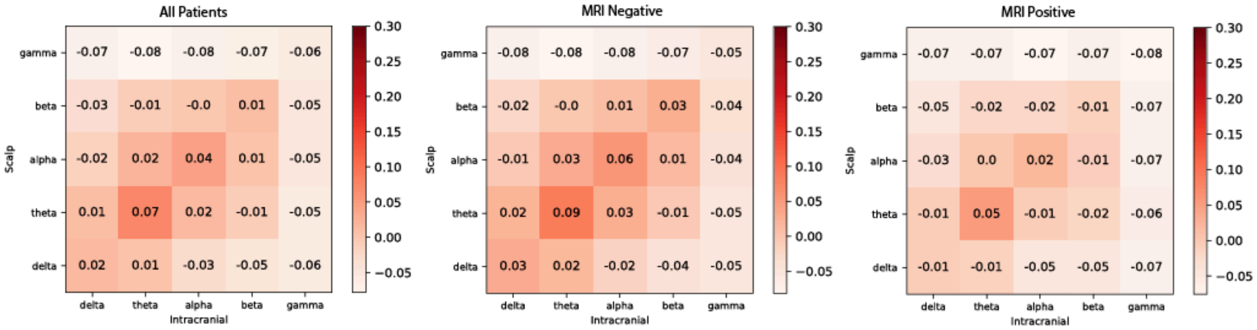


**Figure S3: Mean $R^2$ heatmap for left) all patients, all contacts, center) for all patients with MRI negative (n = 14) and, right) all patients with MRI positive (n = 6).**

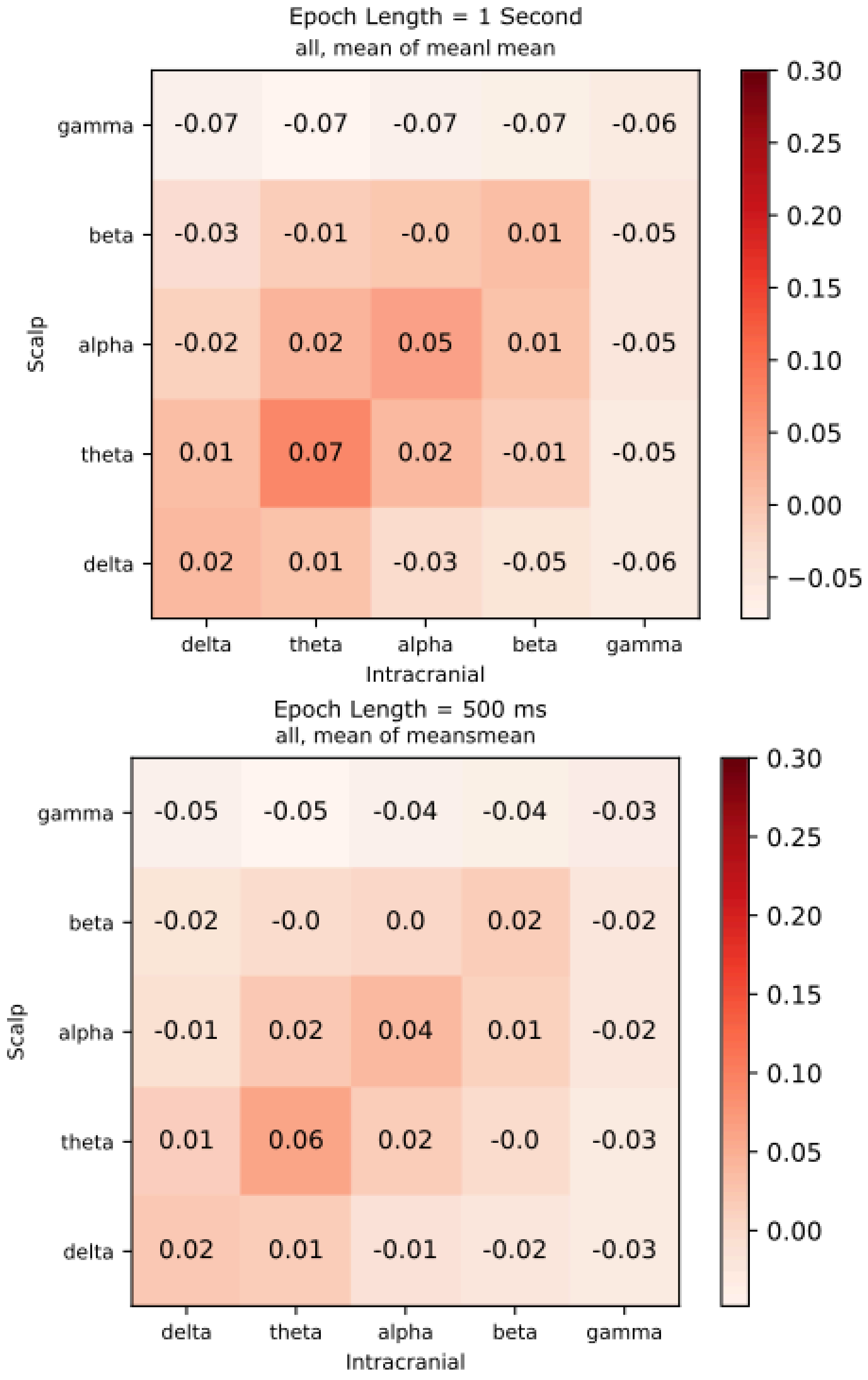


**Figure S4 - Comparison of predictability by bin length**. Plots show mean $\mathbf{R^2}$ of all patients for data that was binned by 1000 millisecond (1 second) and by 500 millisecond epochs. Trends stay similar between both data sets, with a small increase in beta power predictability and small decrease in theta power predictability.

| Region | Subregion | Frequency Band | $R^2$ | Number of Patients | Number of Predictable Electrodes |
|---|---|---|---|---|---|
| Parietal | supramarginal | δ | 0.09 | 1 | 1 |
| | | θ | 0.23 | 8 | 19 |
| | | α | 0.24 | 9 | 23 |
| | | β | 0.22 | 7 | 26 |
| | | γ | 0 | 0 | 0 |
| | precuneus | δ | 0 | 0 | 0 |
| | | θ | 0.23 | 2 | 4 |
| | | α | 0.17 | 2 | 9 |
| | | β | 0.11 | 1 | 3 |
| | | γ | 0 | 0 | 0 |
| | superiorparietal | δ | 0.27 | 4 | 15 |
| | | θ | 0.3 | 6 | 28 |
| | | α | 0.22 | 9 | 31 |
| | | β | 0.23 | 8 | 24 |
| | | γ | 0.15 | 2 | 2 |
| | postcentral | δ | 0.21 | 6 | 14 |
| | | θ | 0.26 | 7 | 20 |
| | | α | 0.26 | 6 | 22 |
| | | β | 0.35 | 5 | 19 |
| | | γ | 0.3 | 1 | 2 |
| | inferiorparietal | δ | 0.16 | 5 | 7 |
| | | θ | 0.2 | 7 | 32 |
| | | α | 0.25 | 9 | 40 |
| | | β | 0.21 | 6 | 33 |
| | | γ | 0.09 | 1 | 3 |
| Put | Put | δ | 0 | 0 | 0 |
| | | θ | 0.15 | 3 | 3 |
| | | α | 0.29 | 3 | 3 |
| | | β | 0.45 | 2 | 2 |
| | | γ | 0 | 0 | 0 |
| Hip | Hip | δ | 0.15 | 4 | 7 |
| | | θ | 0.28 | 3 | 13 |
| | | α | 0.15 | 5 | 10 |
| | | β | 0.11 | 2 | 4 |
| | | γ | 0.09 | 1 | 2 |
| Occipital | pericalcarine | δ | 0 | 0 | 0 |
| | | θ | 0.11 | 1 | 1 |
| | | α | 0.14 | 2 | 2 |
| | | β | 0 | 0 | 0 |
| | | γ | 0 | 0 | 0 |
| | lateraloccipital | δ | 0.17 | 7 | 28 |
| | | θ | 0.19 | 14 | 89 |
| | | α | 0.2 | 12 | 94 |
| | | β | 0.16 | 11 | 69 |
| | | γ | 0.11 | 2 | 4 |
| | cuneus | δ | 0 | 0 | 0 |
| | | θ | 0 | 0 | 0 |
| | | α | 0.23 | 1 | 2 |
| | | β | 0.14 | 2 | 2 |
| | | γ | 0 | 0 | 0 |
| | lingual | δ | 0.15 | 2 | 2 |
| | | θ | 0.15 | 4 | 6 |
| | | α | 0.18 | 5 | 7 |
| | | β | 0.18 | 5 | 6 |
| | | γ | 0 | 0 | 0 |
| Inf | Inf-Lat-Vent | δ | 0 | 0 | 0 |
| | | θ | 0 | 0 | 0 |
| | | α | 0 | 0 | 0 |
| | | β | 0 | 0 | 0 |
| | | γ | 0 | 0 | 0 |

| Region | Subregion | Frequency Band | $R^2$ | Number of Patients | Number of Predictable Electrodes |
|---|---|---|---|---|---|
| Frontal | parsorbitalis | δ | 0.24 | 5 | 29 |
| | | θ | 0.31 | 5 | 30 |
| | | α | 0.28 | 6 | 29 |
| | | β | 0.23 | 5 | 9 |
| | | γ | 0.29 | 1 | 2 |
| | frontalpole | δ | 0.3 | 4 | 31 |
| | | θ | 0.35 | 3 | 31 |
| | | α | 0.21 | 3 | 24 |
| | | β | 0.2 | 2 | 11 |
| | | γ | 0.14 | 1 | 1 |
| | medialorbitofrontal | δ | 0.3 | 3 | 9 |
| | | θ | 0.39 | 3 | 10 |
| | | α | 0.21 | 3 | 8 |
| | | β | 0.1 | 1 | 2 |
| | | γ | 0.11 | 1 | 1 |
| | superiorfrontal | δ | 0.27 | 3 | 9 |
| | | θ | 0.28 | 4 | 13 |
| | | α | 0.22 | 3 | 11 |
| | | β | 0.16 | 2 | 6 |
| | | γ | 0 | 0 | 0 |
| | lateralorbitofrontal | δ | 0.32 | 3 | 14 |
| | | θ | 0.39 | 3 | 16 |
| | | α | 0.17 | 4 | 11 |
| | | β | 0.13 | 3 | 5 |
| | | γ | 0.14 | 2 | 3 |
| | precentral | δ | 0.16 | 2 | 2 |
| | | θ | 0.3 | 3 | 6 |
| | | α | 0.24 | 4 | 7 |
| | | β | 0.39 | 3 | 5 |
| | | γ | 0 | 0 | 0 |
| | parsopercularis | δ | 0.26 | 3 | 9 |
| | | θ | 0.37 | 3 | 14 |
| | | α | 0.2 | 3 | 11 |
| | | β | 0.14 | 2 | 7 |
| | | γ | 0.15 | 1 | 1 |
| | | δ | 0.57 | 1 | 2 |
| | | θ | 0.28 | 3 | 7 |
| | | α | 0.22 | 3 | 5 |
| | | β | 0.19 | 3 | 4 |
| | | γ | 0.22 | 1 | 2 |
| | caudalmiddlefrontal | δ | 0.3 | 4 | 22 |
| | | θ | 0.32 | 6 | 31 |
| | | α | 0.24 | 5 | 23 |
| | | β | 0.31 | 4 | 22 |
| | | γ | 0.4 | 1 | 3 |
| | | δ | 0.33 | 2 | 18 |
| | | θ | 0.37 | 2 | 20 |
| | | α | 0.22 | 2 | 13 |
| | | β | 0.21 | 2 | 10 |
| | | γ | 0 | 0 | 0 |
| | paracentral | δ | 0.2 | 2 | 3 |
| | | θ | 0.32 | 3 | 5 |
| | | α | 0.18 | 1 | 4 |
| | | β | 0.24 | 1 | 3 |
| | | γ | 0 | 0 | 0 |
| Insula | insula | δ | 0.2 | 7 | 12 |
| | | θ | 0.3 | 10 | 28 |
| | | α | 0.2 | 10 | 21 |
| | | β | 0.25 | 5 | 9 |
| | | γ | 0.14 | 2 | 3 |
| Amy | Amy | δ | 0 | 0 | 0 |
| | | θ | 0.1 | 1 | 2 |
| | | α | 0 | 0 | 0 |
| | | β | 0 | 0 | 0 |
| | | γ | 0 | 0 | 0 |

| Region | Subregion | Frequency Band | $R^2$ | Number of Patients | Number of Predictable Electrodes |
|---|---|---|---|---|---|
| Temporal | bankssts | δ | 0.161 | 3 | 5 |
| | | θ | 0.413 | 3 | 6 |
| | | α | 0.222 | 5 | 7 |
| | | β | 0.206 | 4 | 8 |
| | | γ | 0 | 0 | 0 |
| | parahippocampal | δ | 0.131 | 3 | 3 |
| | | θ | 0.328 | 2 | 2 |
| | | α | 0.193 | 2 | 3 |
| | | β | 0.165 | 1 | 2 |
| | | γ | 0.107 | 2 | 2 |
| | entorhinal | δ | 0 | 0 | 0 |
| | | θ | 0 | 0 | 0 |
| | | α | 0 | 0 | 0 |
| | | β | 0 | 0 | 0 |
| | | γ | 0 | 0 | 0 |
| | middletemporal | δ | 0.242 | 10 | 43 |
| | | θ | 0.283 | 11 | 59 |
| | | α | 0.187 | 11 | 42 |
| | | β | 0.15 | 8 | 27 |
| | | γ | 0.278 | 1 | 1 |
| | inferiortemporal | δ | 0.25 | 7 | 16 |
| | | θ | 0.302 | 6 | 18 |
| | | α | 0.223 | 6 | 12 |
| | | β | 0.153 | 3 | 7 |
| | | γ | 0 | 0 | 0 |
| | temporalpole | δ | 0 | 0 | 0 |
| | | θ | 0.229 | 2 | 4 |
| | | α | 0.095 | 1 | 3 |
| | | β | 0 | 0 | 0 |
| | | γ | 0 | 0 | 0 |
| | fusiform | δ | 0.137 | 3 | 7 |
| | | θ | 0.271 | 4 | 12 |
| | | α | 0.211 | 5 | 12 |
| | | β | 0.14 | 3 | 4 |
| | | γ | 0.163 | 1 | 1 |
| | transversetemporal | δ | 0 | 0 | 0 |
| | | θ | 0 | 0 | 0 |
| | | α | 0 | 0 | 0 |
| | | β | 0 | 0 | 0 |
| | | γ | 0.082 | 1 | 1 |
| | superiortemporal | δ | 0.157 | 3 | 3 |
| | | θ | 0.236 | 6 | 8 |
| | | α | 0.177 | 4 | 13 |
| | | β | 0.251 | 2 | 5 |
| | | γ | 0 | 0 | 0 |
| Cingulate | isthmuscingulate | δ | 0.16 | 2 | 4 |
| | | θ | 0.289 | 2 | 3 |
| | | α | 0.149 | 2 | 3 |
| | | β | 0.139 | 1 | 2 |
| | | γ | 0 | 0 | 0 |
| | posteriorcingulate | δ | 0.372 | 1 | 3 |
| | | θ | 0.261 | 2 | 4 |
| | | α | 0.187 | 1 | 3 |
| | | β | 0.215 | 1 | 3 |
| | | γ | 0 | 0 | 0 |
| | caudalanteriorcingulate | δ | 0.35 | 2 | 4 |
| | | θ | 0.324 | 2 | 5 |
| | | α | 0.226 | 2 | 4 |
| | | β | 0.112 | 1 | 2 |
| | | γ | 0 | 0 | 0 |
| | rostralanteriorcingulate | δ | 0.221 | 3 | 7 |
| | | θ | 0.361 | 3 | 6 |
| | | α | 0.18 | 3 | 3 |
| | | β | 0 | 0 | 0 |
| | | γ | 0 | 0 | 0 |

**Table S3: Mean like-band regression results across the contact population, divided by brain region.**

| parcellation | # predictable contacts | # total contacts | % predictable |
|---|---|---|---|
| amygdala | 4 | 4 | 100.0% |
| putamen | 4 | 5 | 80.0% |
| rostralanteriorcingulate | 12 | 16 | 75.0% |
| parsopercularis | 15 | 23 | 65.2% |
| medialorbitofrontal | 9 | 14 | 64.3% |
| pericalcarine | 9 | 14 | 64.3% |
| middletemporal | 99 | 169 | 58.6% |
| temporalpole | 9 | 16 | 56.3% |
| paracentral | 9 | 16 | 56.3% |
| frontalpole | 27 | 48 | 56.3% |
| cuneus | 10 | 18 | 55.6% |
| fusiform | 25 | 45 | 55.6% |
| superiortemporal | 45 | 83 | 54.2% |
| lingual | 17 | 32 | 53.1% |
| isthmuscingulate | 9 | 17 | 52.9% |
| rostralmiddlefrontal | 21 | 40 | 52.5% |
| precentral | 19 | 38 | 50.0% |
| insula | 49 | 100 | 49.0% |
| inferiortemporal | 38 | 78 | 48.7% |
| lateraloccipital | 154 | 318 | 48.4% |
| inferiorparietal | 74 | 159 | 46.5% |
| superiorparietal | 34 | 82 | 41.5% |
| superiorfrontal | 12 | 29 | 41.4% |
| parsorbitalis | 28 | 69 | 40.6% |
| bankssts | 16 | 40 | 40.0% |
| transversetemporal | 8 | 20 | 40.0% |
| supramarginal | 35 | 88 | 39.8% |
| postcentral | 29 | 73 | 39.7% |
| caudalmiddlefrontal | 23 | 58 | 39.7% |
| parahippocampal | 7 | 19 | 36.8% |
| lateralorbitofrontal | 13 | 37 | 35.1% |
| hippocampus | 22 | 67 | 32.8% |
| precuneus | 9 | 28 | 32.1% |
| posteriorcingulate | 3 | 10 | 30.0% |
| entorhinal | 2 | 7 | 28.6% |
| caudalanteriorcingulate | 3 | 12 | 25.0% |
| parstriangularis | 4 | 25 | 16.0% |

**Table S4 Proportion of contacts in each region that are predictable.** Sorted in descending order by percent predictable. All regions contain at least 1 contact from the total sample size of 1918 contacts that were predictable.

| Region | Frequency Band | Degrees Freedom | T Statistic | P Value |
|---|---|---|---|---|
| Occipital | Delta | 16 | 1.199 | 0.24 |
| Occipital | Theta | 16 | 0.203 | 0.84 |
| Occipital | Alpha | 16 | -0.14 | 0.89 |
| Occipital | Beta | 16 | -1.884 | 0.078 |
| Occipital | Gamma | 16 | 1.975 | 0.066 |
| Frontal | Delta | 14 | -2.314 | 0.036 |
| Frontal | Theta | 14 | 0.439 | 0.667 |
| Frontal | Alpha | 14 | -2.23 | 0.043 |
| Frontal | Beta | 14 | -2.904 | 0.011 |
| Frontal | Gamma | 14 | -1.734 | 0.104 |
| Temporal | Delta | 18 | 0.552 | 0.587 |
| Temporal | Theta | 18 | -0.325 | 0.73 |
| Temporal | Alpha | 18 | -0.845 | 0.409 |
| Temporal | Beta | 18 | -1.315 | 0.205 |
| Temporal | Gamma | 18 | -0.541 | 0.594 |
| Parietal | Delta | 19 | 0.567 | 0.577 |
| Parietal | Theta | 19 | -0.186 | 0.855 |
| Parietal | Alpha | 19 | 0.939 | 0.359 |
| Parietal | Beta | 19 | -0.476 | 0.639 |
| Parietal | Gamma | 19 | 1.352 | 0.192 |

**Table S5: One-Sample T-Test results for patient regional slope averages of predictability vs. distance away from scalp.**

| Brain Region | Intracranial Band | Scalp EEG Band | | | |
|---|---|---|---|---|---|
| | | Delta | Theta | Alpha | Beta |
| Prefrontal Cortex | Beta | — | — | — | 0.07 * |
| | Alpha | 0.08 * | 0.09 * | 0.07 * | — |
| | Theta | 0.11 | 0.15 * | — | — |
| | Delta | — | — | — | — |
| Mesial Temporal Lobe | Beta | 0.03 * | — | — | — |
| | Alpha | 0.04 * | 0.06 * | 0.03 * | — |
| | Theta | 0.07 * | 0.10 *** | — | — |
| | Delta | 0.07 * | 0.04 * | — | — |
| Orbitofrontal Cortex | Beta | — | — | — | 0.02 * |
| | Alpha | 0.05 * | 0.06 * | 0.04 * | — |
| | Theta | 0.09 | 0.11 * | — | — |
| | Delta | — | — | — | — |

$R^2$ Value Scale

0.00 0.05 0.10 0.15 0.20

* p < 0.05, ** p < 0.01, *** p < 0.001, — = not significant

**Table S6: Significant regressions in regional Leave-one-patient-out CV models.** Significance was tested with one-sample T-tests, with null hypothesis mean of all left-out-patients' predictability of having R 0

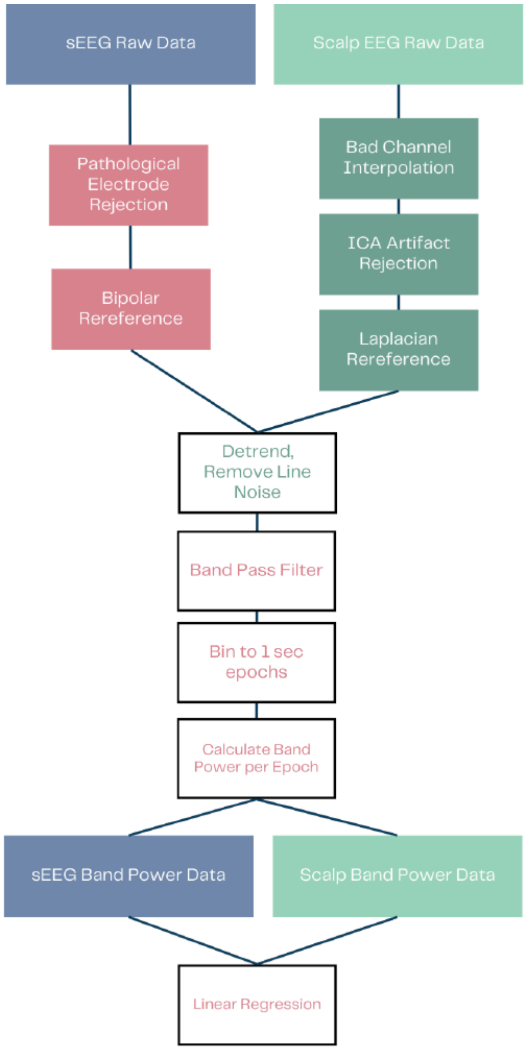


**Figure S5: Flowchart of electrophysiology preprocessing for scalp EEG and intracranial EEG (StereoEEG or sEEG).**

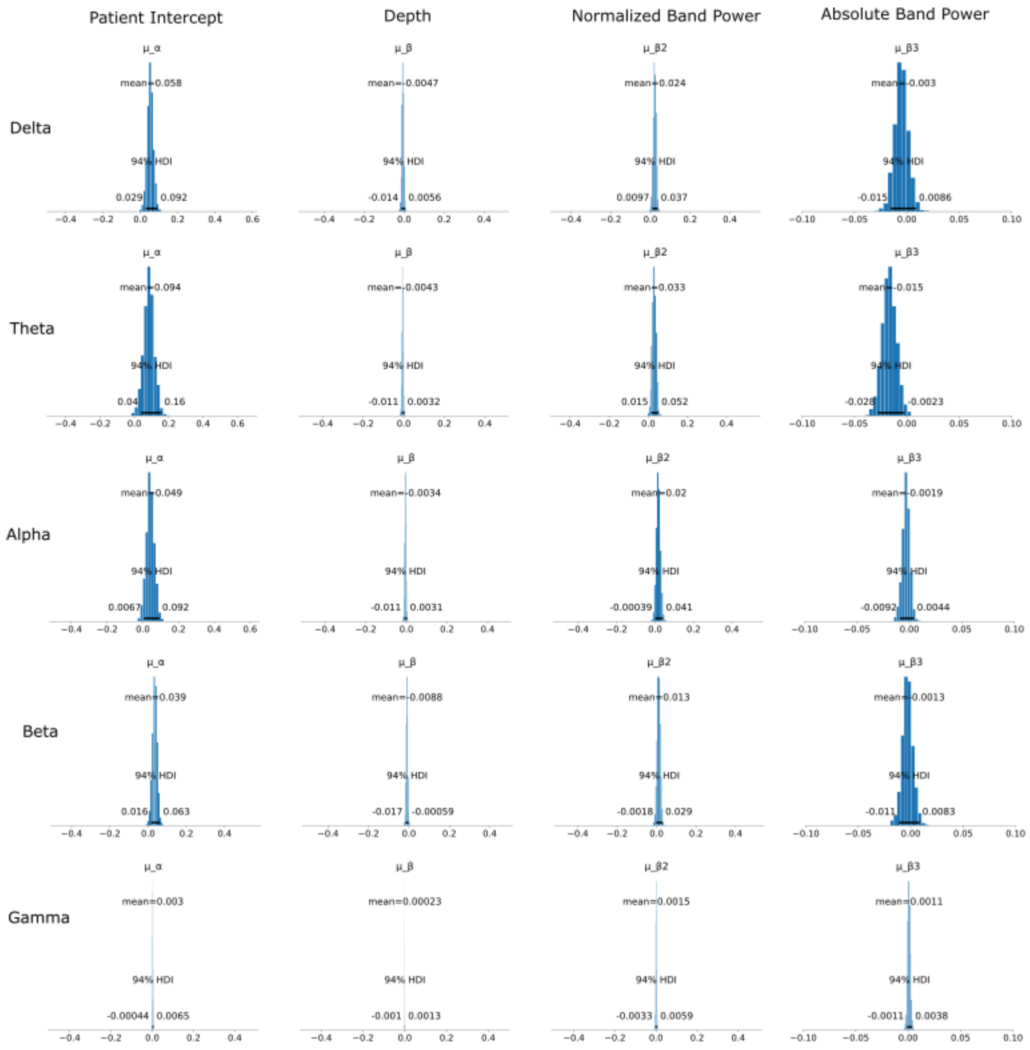


**Figure S6: Mean posterior distributions of each of the 3 beta coefficient parameters and alpha in the hierarchical Bayesian model of contact predictability across each of the like-band regressions.**

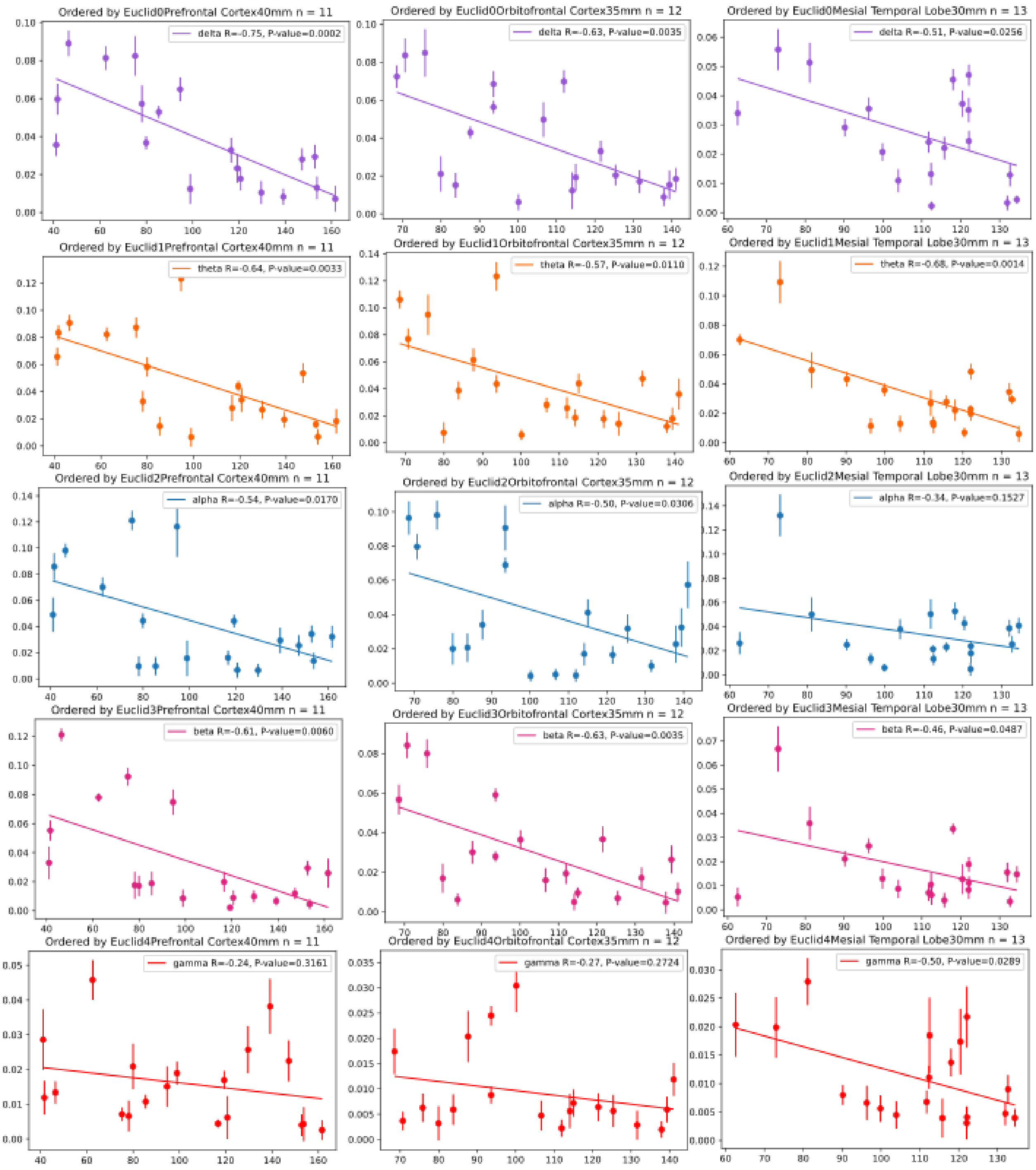


**Figure S7: Visualization of mean beta coefficients, representing the weighting of scalp electrodes, in Leave-One-Patient-Out regression**

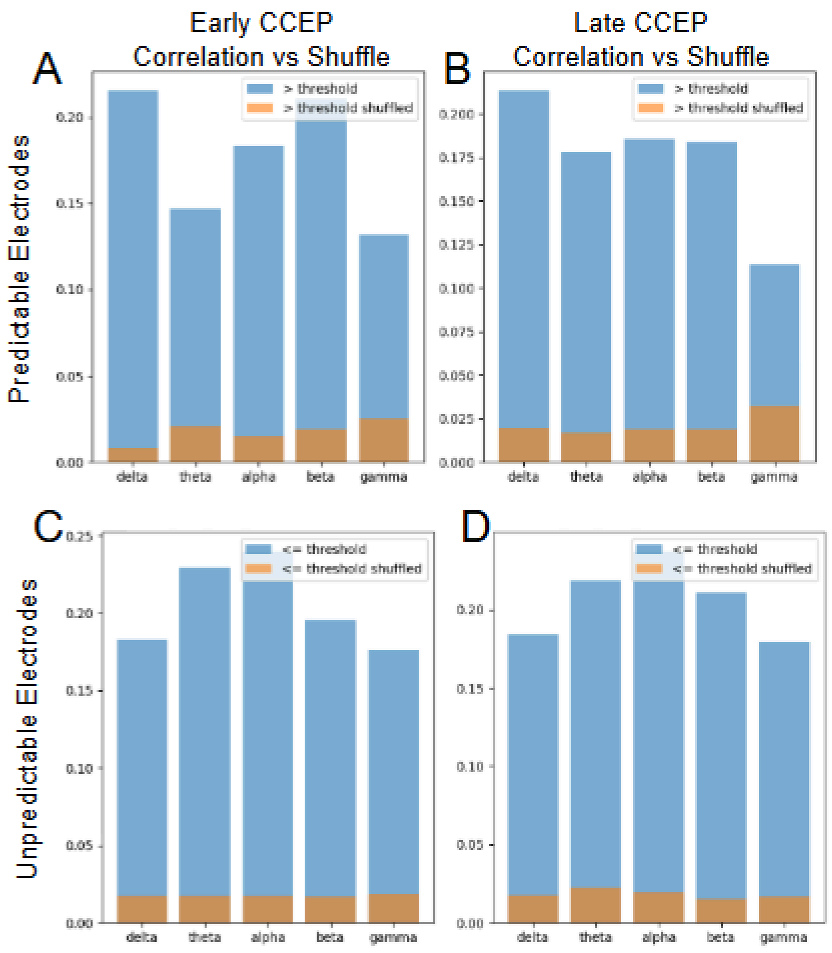


**Figure S8. Correlation between spontaneous prediction and evoked responses across frequency bands.** Bar graphs comparing predictable versus unpredictable contacts across five frequency bands (delta, theta, alpha, beta, gamma). **(A)** Early CCEP (Blue) correlation versus shuffled data (Orange) correlation within predictable contacts ($R^2 > 0$). **(B)** Late CCEP correlation versus shuffled data. **(C, D)** Repeat analysis but with unpredictable contacts ($R^2 <= 0$).

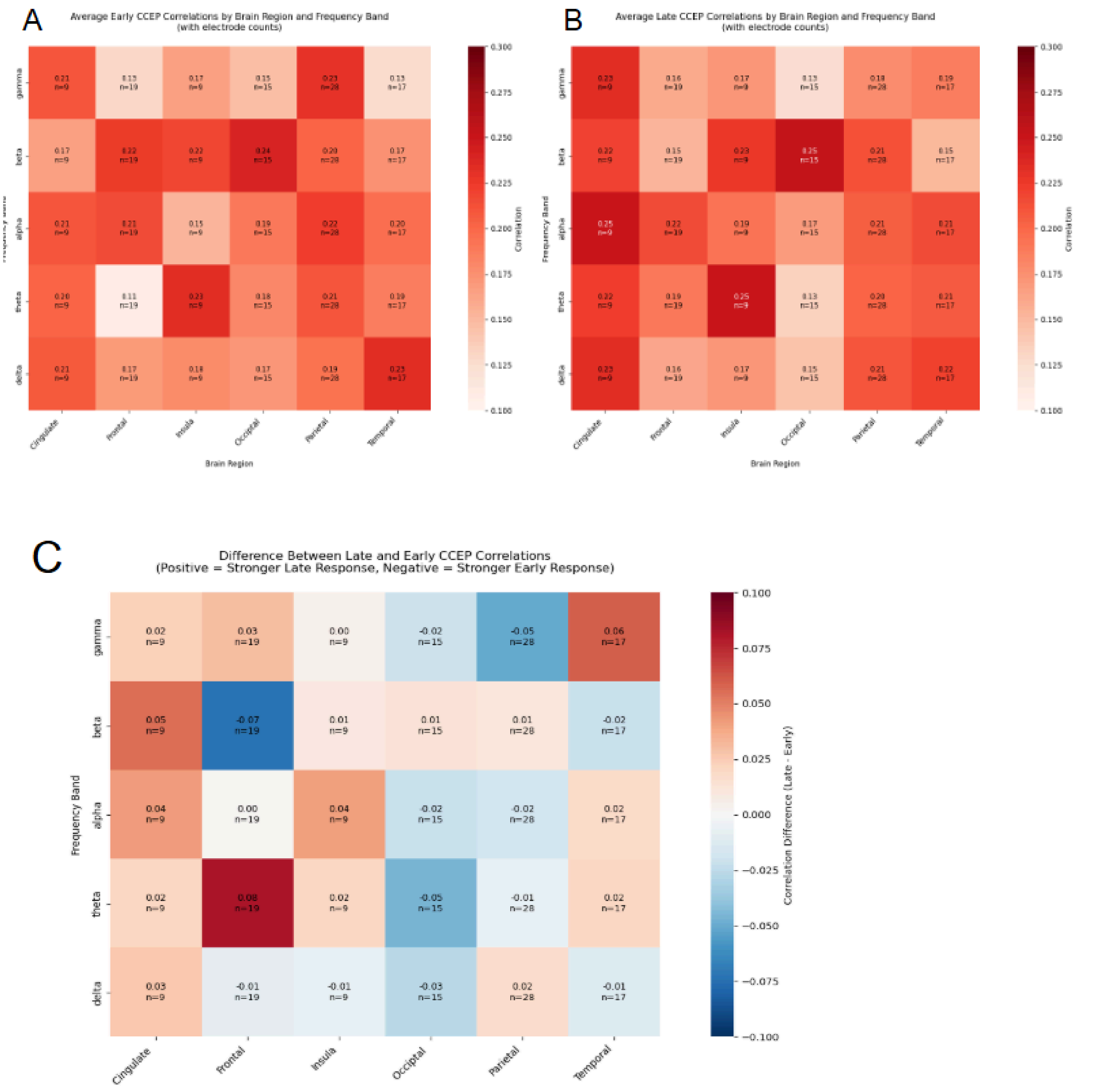


**Figure S9 - Regional analysis showing the anatomically-specific difference** in correlations between spontaneous predictive maps and early and late sCCEP, with strongest differences found in cingulate and insula (theta band) and lateral temporal regions (alpha band). All correlations significantly exceeded chance levels based on shuffling analyses ($p<0.001$). (A) Early CCEP correlations by brain region and frequency band. Warmer colors indicate stronger correlations (B) Late CCEP correlations (C) Difference between Matrix A and B, showing regions that tend to have differing correlations to early vs late CCEP. Positive and warmer colors indicate a stronger late response than early, and negative and colder colors indicate the opposite
